# Structure of Full Length Plasmodium Myosin A and its light chain PfELC, dual targets against malaria parasite pathogenesis

**DOI:** 10.1101/2020.06.30.179788

**Authors:** Dihia Moussaoui, James P. Robblee, Daniel Auguin, Elena B. Krementsova, Silvia Haase, Thomas C. A. Blake, Jake Baum, Julien Robert-Paganin, Kathleen M. Trybus, Anne Houdusse

**Affiliations:** Structural Motility, Institut Curie, Paris Université Sciences et Lettres, Sorbonne Université, CNRS UMR144, 75005 Paris, France; Department of Molecular Physiology and Biophysics, University of Vermont, USA; Laboratoire de Biologie des Ligneux et des Grandes Cultures (LBLGC), Université d’Orléans, INRA, USC1328, 45067 Orléans, France; Department of Life Sciences, Imperial College London, South Kensington, London SW7 2AZ, UK

## Abstract

Parasites from the genus *Plasmodium* are the causative agents of malaria. The mobility, infectivity and ultimately pathogenesis of this parasite relies on a macromolecular complex, called the glideosome. At the core of the glideosome is an essential and divergent Myosin A motor (PfMyoA), a first order drug target against malaria. Here we present the full-length structure of PfMyoA in two states of its motor cycle. We report novel interactions that are essential for motor priming and the mode of recognition of its two light chains (PfELC and MTIP) by two degenerate IQ motifs. Kinetic and motility assays using PfMyoA variants, along with molecular dynamics, demonstrate how specific priming and atypical sequence adaptations tune the motor’s mechano-chemical properties. Supported by evidence for an essential role of the PfELC in malaria pathogenesis, these structures provide a blueprint for the design of future antimalarials targeting both the glideosome motor and its regulatory elements.

**Highlights:**

- The first structures of the full length PfMyoA motor in two states of its motor cycle.
- A unique priming of the PfMyoA lever arm results from specific lever arm/motor domain interactions, which allows for a larger powerstroke to enhance speed.
- Sequence adaptations within the motor domain and degenerate IQ motifs in the lever arm dictate PfMyoA motor properties.
- PfELC is essential for blood cell invasion and is a weak link in the assembly of a fully functional motor, providing a second novel target for antimalarial drug design.

---

Parasites from the genus *Plasmodium*, the causative agents of malaria, are responsible for half of a million deaths per year^1^. Significant effort and money has been devoted to developing vaccines and new preventive treatments because the malaria parasites are becoming resistant to current artemisinin-based therapies^2^. The global death rate from malaria has recently started to rise after many years of decrease^1^, emphasizing the necessity to develop new interventions, particularly because climate change may further expand the range of *Anopheles* mosquitoes.

The mosquito-borne parasite *P. falciparum*, the deadliest species that infects humans, alternates between motile and non-motile stages. Locomotion of apicomplexan parasites occurs by a process called gliding motility (reviewed in^3^). This mode of displacement as well as the infectivity of the parasite relies on a macromolecular assembly called the glideosome that is anchored in an inner membrane complex located ^~^25 nm below the parasite plasma membrane. The core of the glideosome consists of an atypical class XIV myosin A (PfMyoA) and a divergent actin (PfAct1). PfMyoA is a short myosin with a conserved globular motor domain and a lever arm that binds two light-chains: an essential light chain (PfELC) and MTIP (<u>m</u>yosin <u>t</u>ail <u>i</u>nteracting <u>p</u>rotein)^4,5^. The N-terminal extension of MTIP binds to integral membrane proteins called GAPs (<u>g</u>lideosome <u>a</u>ssociated <u>p</u>roteins), which anchors the myosin so that it can interact cyclically with actin^6^ (**Fig. 1a**) (reviewed in^3^). PfMyoA is a critical molecule in the parasite life-cycle because it powers the fast motility (^~^2 μm.s^−1^) required during motile sporozoite stages^7^, and is essential for providing the force (up to 40 pN) needed for non-motile merozoites to invade erythrocytes^8,9^.

**Figure 1 –.**
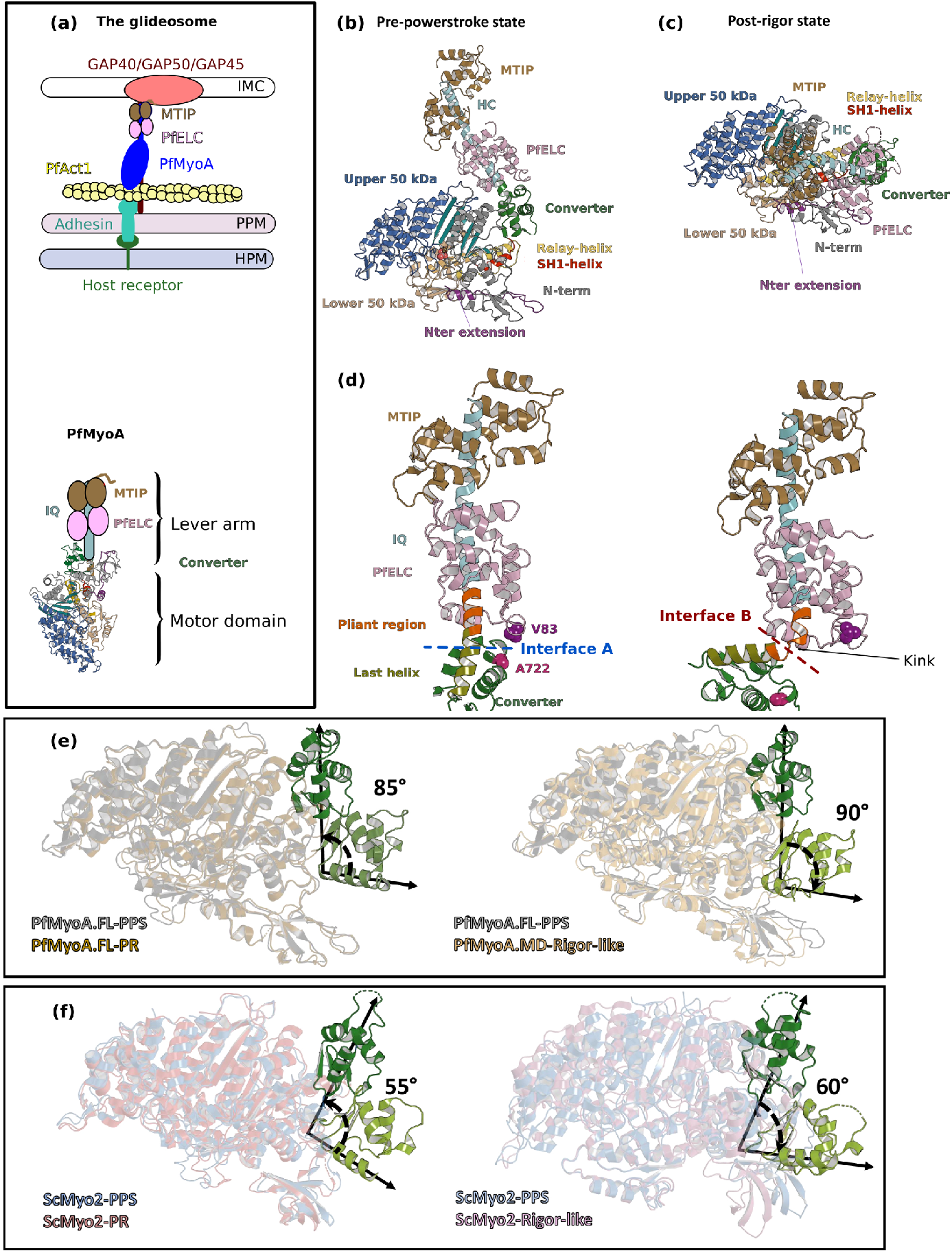
Crystal structures of PfMyoA in the Post-rigor (PR) and the Pre-Powerstroke (PPS) states. **(a) (Left)** The PfMyoA motor is located in the intermembrane space of the parasite. PfMyoA (blue) binds two light-chains, PfELC and MTIP. MTIP connects the motor to the glideosome-associated proteins (GAP) complex, which is anchored in the Inner Membrane Complex (IMC). PfMyoA cyclically interacts with PfAct1 filaments, which are bound to adhesins from the Parasite Plasma Membrane (PPM); these adhesins also bind receptors from the Host cell Plasma Membrane (HPM). The displacement of PfAct1 filaments by PfMyoA drives parasite gliding motility. **(Right)** The crystal structure of the motor domain of PfMyoA has been solved^9^, but the lever arm structure was not known. **(b,c)** Overall structures of the full length PfMyoA motor in the PPS and PR states, displayed so that their N-terminal subdomains adopt a similar orientation. As expected, the orientation of the Converter and lever arm differs in these two states. **(d)** The lever arm has been built in these two states of the motor, revealing the structure of the two bound light chains, PfELC and MTIP, displayed here in a similar orientation. The kink in the lever arm helix at the end of the Converter (pliant region in orange; last helix of the Converter in deep olive green) induces different Converter/PfELC interfaces in the PPS (interface A, left) compared to the PR state (interface B, right). To illustrate that the two interfaces are different, two reporter residues are displayed as spheres, A722 from the Converter and V83 from PfELC. These residues are part of Interface A but not part of Interface B. **(e)** and **(f)** represents the recovery stroke (left) and the powerstroke (right) for PfMyoA and Scallop myosin 2 respectively. Structures of scallop myosin 2 used: PR (PDB code 1S5G); PPS (PDB code 1QVI).

The molecular motor myosin has a conserved ATPase cycle in which the state of hydrolysis of the nucleotide drives both the association with actin and movements of the lever arm. Structural reorganization initiated at the active site are transmitted allosterically by connectors, and amplified by the lever arm (**Supplementary Fig. 1a, 1c, 1d**) (reviewed by^10^). We recently solved the X-ray structures of the *P. falciparum* MyoA motor domain in three states of this ATPase cycle^9^, which revealed how the lack of consensus residues in the connectors are compensated by the presence of a unique N-terminal extension of the heavy chain (**Supplementary Fig. 1c**). Phosphorylation of Ser19 within this extension directly tunes speed and force production^9^. The motor moves actin at high-speed and exerts low ensemble force when phosphorylated; conversely, it produces more force at the expense of slower speed when unphosphorylated^9^ (**Supplementary Fig. 1b**). According to this model, phosphorylated PfMyoA would allow the parasite to move at high velocity during motile stages such as the sporozoites, and when unphosphorylated, would provide the high force necessary for merozoites to enter erythrocytes in invasive stages^9^.

The motor domain is not the only region of PfMyoA lacking myosin consensus sequences. The lever arm typically contains “IQ motifs” (consensus sequence IQxxxRGxxxR) that bind light chains or calmodulin. In PfMyoA, both the first IQ motif and the PfELC that binds to it are so degenerate in their sequence that the existence of an essential light chain was only recently recognized^4,5^ (**Fig. 1a**). The structure of MTIP in complex with a *Plasmodium* IQ2 peptide is the only structural information available for the PfMyoA lever arm^11,12,13^. Further structures are required to establish how interactions between the motor domain and the lever arm influence the overall dynamics and properties of the motor.

Here we present the X-ray structures of full-length PfMyoA complexed to PfELC and MTIP in two states of the motor cycle. These structures, together with molecular dynamics and small angle X-ray scattering (SAXS), reveal the specific orientation of the lever arm in the pre-powerstroke state. This atypical priming, which enables a larger powerstroke, is stabilized by the Converter and PfELC forming specific interactions with the motor domain. Kinetic and motility assays on mutant PfMyoA constructs show how the specific priming and unique sequence adaptations tune the motor properties of this atypical myosin. The lever arm structure explains how the PfELC and MTIP light chains evolved to recognize degenerate IQ motifs. The PfELC, a weak link in forming a fully functional motor, is shown here to be essential for red blood cell invasion, thus providing a novel target for the design of anti-malarial compounds.

## Results

### Full-length structures of PfMyoA reveal a specific priming of the lever arm

We determined the crystal structures of Full-length PfMyoA with two light chains bound (PfMyoA/PfELC/MTIP-Δn) in the post-rigor (PfMyoA•FL-PR) and in the pre-powerstroke (PfMyoA•FL-PPS) states at 2.5 Å and 3.9 Å resolution, respectively (**Supplementary Table 1**) (**Fig. 1b, 1c**). The structure of a similar construct in which the N-terminal HC extension (Nter) of the PfMyoA heavy chain (HC) was truncated (PfMyoA-ΔNter/PfELC/MTIP-Δn) was also solved in the Post-Rigor (PR) state at 3.3 Å resolution (PfMyoA•ΔNter-PR) (**Supplementary Fig. 2a, 2b**). For all three structures, the electron density was well-defined for the motor domain and the lever arm, in particular for the interfaces between the HC and the PfELC and the interface between the two light-chains (**Supplementary Fig. 3**).

These structures allow the description of the lever arm that was not present in the previously published PfMyoA motor domain (MD) structures^9^. When comparing the PR and PPS states, the overall fold of both the IQ region of the HC/PfELC/MTIP and Converter regions are highly similar. Major differences are located at the ELC/Converter interface when the PR and the PPS are compared (**Fig. 1d**), predominantly due to a sharp kink in the pliant region^14^ observed in the PR structures that promotes contacts between the ELC and the motor domain (**Supplementary Fig. 2c**). SAXS experiments demonstrate that this ‘folded-back’ position of the lever arm is not primarily populated in solution and is selected by crystal contacts. In solution, the lever arm adopts an extended conformation as seen in other myosins (**Supplementary Data 1; Supplementary Fig. 2d, 2e-h**).

The Converter orientation is the same in both the motor domain (PfMyoA•MD-PR)^9^ and the fulllength (PfMyoA•FL-PR) post-rigor structures (**Supplementary Fig. 4a**). In the PPS states crystallized with ADP and Pi analogs, however, the Converter orientation is less primed by ^~^30° in the MD structure compared with that found in the FL structure^9^ (**Supplementary Fig. 4b**). SAXS experiments validated that the PPS of PfMyoA adopts the more primed orientation in solution (**Supplementary Data 1; Supplementary Fig. 2e**). The MD structure with ADP.VO4 bound thus likely corresponds to an intermediate state populated during the recovery stroke, close to that of the fully primed PPS (**Supplementary Fig. 4c, 4d**). Interestingly, the priming of the lever arm in the TgMyoA•MD-PPS structure^15^ is identical to that of the PfMyoA•FL-PPS structure (**Supplementary Fig. 4e**). No major difference is found in the motor domain when the FL and MD structures are compared (**Supplementary Fig. 4f**). Notably, the lever arm is more primed in the PPS state of PfMyoA compared with that of conventional myosins, leading to a larger powerstroke than most myosins.

The FL structures of PfMyoA can be used to define the lever arm swing occurring during the PfMyoA motor cycle. The Converter is reoriented ^~^90° during the powerstroke (**Fig. 1e**). In conventional myosins, such as Scallop Myosin 2 (ScMyo2), the Converter is reoriented only ^~^60° during the powerstroke (**Fig. 1f**). In comparison, Myo10 undergoes a large ^~^140° swing during the powerstroke^16^ linked to both a highly primed lever arm orientation in its PPS state and an unusual Rigor Converter position. Specific structural features of PfMyoA thus control the lever arm repriming and larger swing amplitude during the powerstroke.

### Sequence adaptations tune distinct transitions of the cycle

Specific sequence differences in PfMyoA do not alter the global positioning of subdomains compared with conventional myosins, but instead affect the stability of structural states the motor explores and tune the kinetics of transitions between these states. Deletion of the Nter enhanced the duty ratio (time spent strongly bound to actin) by greatly slowing ADP release to the extent that it rate-limited the overall ATPase cycle, which is ^~^14-fold slower than WT^9^. The similarity of the ΔNter heavy chain and WT PfMyoA structures (rmsd of 0.415 Å on 993 atoms) (**Supplementary Fig. 2b**) strongly argues that the dramatic kinetic differences observed between these two constructs are due to an altered equilibrium between the states of the motor cycle^9^.

We previously showed that Serine 19 phosphorylation (SEP19) accelerates ADP release by stabilizing the Converter in its Rigor orientation through a polar interaction with ^Converter^K764 but other residues likely modulate these motor properties. In the Rigor state, ^Nter-extension^E6 establishes cation-π stacking interactions with ^Switch-2^F476 and ^Switch-1^R241 from the Switch-2 and Switch-1 elements of the active site^9^ (**Fig. 2a**). The charge reversal E6R mutant was designed to disrupt the interaction seen in the Rigor state, a state the motor needs to populate in order to release ADP. The effects of the E6R mutation are similar to that observed with mutants disrupting the polar bond between SEP19 and the Converter (S19A and K764E)^9^): ^~^2-fold reduced maximal actin-activated ATPase, ^~^2-fold reduced rate of ADP release that is correlated with a reduced speed of moving actin, ^~^3-fold increased ensemble force, and ^~^2.5-fold faster dissociation of acto-PfMyoA by MgATP (**Fig. 2b-f, Supplementary Table 2**). All these results are consistent with E6 stabilizing the Rigor state in WT via its interactions with Switches-1 and −2.

**Figure 2 –.**
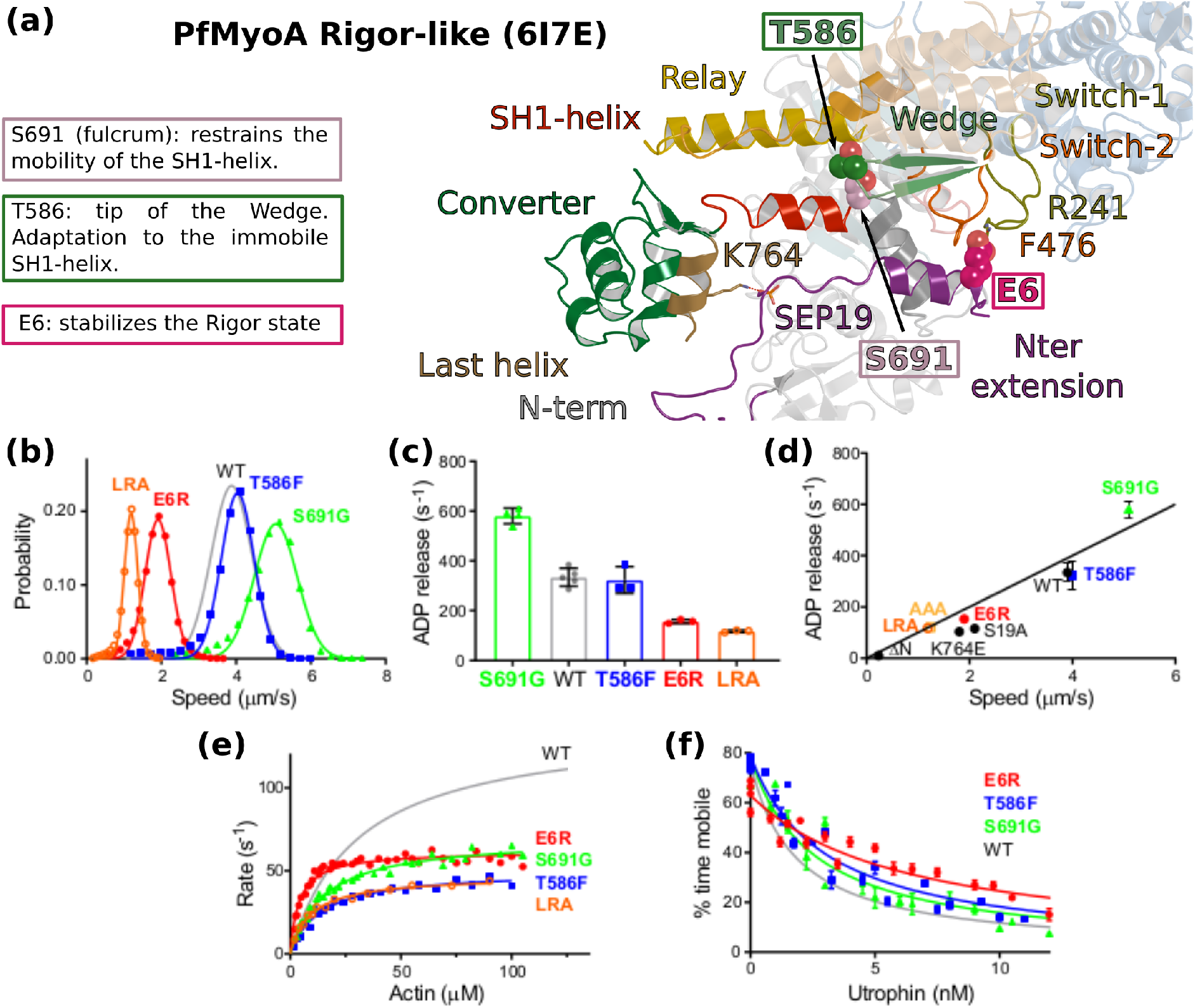
Sequence adaptations tune distinct transitions of the cycle. **(a)** Location and function of three mutated residues. **(b)** Speed distributions from representative in vitro motility assays. **Supplementary Table 2** shows values from additional experiments. **(c)** ADP release rates from acto-PfMyoA. WT data are from^9^. Data for LRA (3 experiments, 2 protein preparations); E6R (3 experiments, 3 protein preparations); T586F (3 experiments, 2 protein preparations); S691G (4 experiments, 3 protein preparations). Values, mean ± SD. **(d)** Correlation of ADP release rates and in vitro motility speed. **(e)** Actin-activated ATPase activity. WT data are from^9^. Data from at least 2 protein preparations and 2 experiments for each construct were fitted to the Michaelis-Menten equation. Error, SE of the fit. **(f)** Ensemble force measurements using a utrophin-based loaded in vitro motility assay. A myosin that produces more force requires higher utrophin concentrations to slow motion: E6R, 4.02 ± 0.31 nM; T586F, 2.38 ± 0.18 nM; S691G, 1.99 ± 0.19 nM; WT, 1.40 ± 0.08 nM. WT data are from^9^. Error, SE of the fit. Data from 2 protein preparations and 3 experiments for each mutant construct. **Supplementary Fig. 8d** shows additional data with an expanded x-axis. Skeletal actin was used for all experiments. Temperature, 30°C. See also **Supplementary Table 2** for values.

Two residues that are conserved in most myosins but not in PfMyoA were mutated to further investigate the role of sequence differences that are unique to PfMyoA (S691G and T586F). These residues are part of the environment between the Relay and SH1 helix, two critical connectors for the control of Converter repriming and the powerstroke (**Fig. 2a**). The presence of S691 at the base of the SH1 helix, which is typically a glycine in all other myosins, restrains the mobility of the SH1 helix. S691G is the only mutant to date that showed faster *in vitro* motility and faster ADP release than WT, implying that it destabilizes the strong ADP state (**Fig. 2b-d, Supplementary Table 2**). The enhanced speed could be attributed to an increased flexibility of the SH1 helix or a decrease in steric hindrance that favors Converter movement, resulting in faster ADP release. The maximal actin-activated ATPase activity was half that of WT and not rate-limited by ADP release (**Fig. 2e, Supplementary Table 2**). S691 is involved in the communication between the Relay and the SH1 helices by maintaining an electrostatic bond with ^Relay^Q494, which is part of the sequence compensation specific to PfMyoA. This bond is formed in the PR and PPS states but must be lost during the powerstroke. The absence of this bond in the mutant could reduce actin-activated ATPase because of altered communication between the connectors that affects the kinetics of the transitions in the motor cycle, introducing a slower step most likely early in the powerstroke. The 9-fold higher basal ATPase activity in the absence of actin (**Supplementary Table 2**) indicates that the pre-powerstroke (PPS) state of the mutant is less stable than WT, while the similar rate of dissociation of acto-PfMyoA by MgATP implies that the stability of the Rigor state and the transition to PR are like WT (**Supplementary Table 2**). The mutant retains the ability to undergo conformational transitions when bound to actin under strain, with a modestly enhanced ensemble force (1.4-fold) compared with WT (**Fig. 2f, Supplementary Table 2**). The role Ser691 plays in PfMyoA is thus to stabilize the PPS state and to slow the steps that are involved in rearrangements on F-actin to control the powerstroke.

Thr586 from the Wedge was replaced by the bulkier aromatic residue Phe that is present in conventional myosins (T586F). The Wedge guides the movement of the Relay-helix and the SH1-helix connectors depending on L50 subdomain movements that result from events in the active site (**Supplementary Fig. 1c)**. This mutation had relatively little effect on *in vitro* motility speed or on ADP release rates compared with WT (**Fig. 2 b-d, Supplementary Table 2**), despite the fact that the bulkier Phe would likely reduce the mobility around the Wedge due to steric hindrance and thus impede the communication between the active site and the lever arm during the lever arm swing. The rate-limiting step for motility is not the first transition of the powerstroke that leads to the Strong-ADP state on actin, but the following ADP release step, which is not affected by this mutation. Rearrangements near the Wedge thus play a minor role for ADP release, consistent with previous structural results^17^. The ^~^40% faster transition to the PR state suggests that Rigor is slightly destabilized, while the elevated basal ATPase indicates a reduced stability of the PPS state (**Supplementary Table 2**). The reduced maximal actin-activated ATPase activity of the mutant indicates that other transitions in the motor cycle have slowed (**Fig. 2e, Supplementary Table 2**). Similar to the other two point mutants (E6R and S691G), its ability to generate motion under force was better than WT, implying more time spent in actin-bound states of the powerstroke (**Fig. 2f, Supplementary Table 2**). Thr586, which is an adaptation to the immobile SH1 helix, contributes to stabilizing both the PPS and Rigor states.

The full-length structures are consistent with the model previously proposed for the atypical and tunable mechanism of force-production^9^. The new mutants analyzed here validate the idea that the sequence compensations observed in the SH1-helix, the Wedge and the Nter are key-elements that not only maintain the allosteric communication between the different myosin connectors, but also tune the motor properties.

### Specific recognition of the degenerate IQ motifs in the atypical PfMyoA Lever Arm

PfMyoA, like other class XIV myosins, is short and lacks a tail domain^18^. The C-terminal region of PfMyoA consists of two degenerate IQ motifs (PfIQ1 and PfIQ2) that deviate from the consensus IQ sequence (IQxxxRGxxxR) but are recognized by the native light-chains, PfELC and MTIP. The recognition of PfIQ2 by MTIP has already been described^11,12,13^ and involves several residues from the consensus IQ motif sequence (**Supplementary data 2, Supplementary Fig. 5 a-e**).

The FL structures allow the description of how PfELC binds to the lever arm and the interactions it forms with the motor domain. PfELC interacts with the long HC α-helix with its N-lobe in a closed conformation and its C-lobe in a semi-open conformation, as it is the case for other ELCs (**Fig. 3**)^19^. While the structure of the N-lobe is mostly conserved compared to other ELCs (**Fig. 3a, 3b**)^20^, the C-lobe contains an extremely short α5 helix which consists of only one turn (α5*) followed by a short linker in which a small inserted helix turn forms (α5’*) (**Fig. 3c, 3d**). In addition, the inter-lobe linker is shorter than canonical ELCs and forms an internal hairpin-like structure, that interacts strongly with the N-lobe and does not contact residues of the HC helix (**Fig. 3e, 3f**). This unusual feature for an inter-lobe linker prevents the PfELC from fully surrounding the HC helix.

**Figure 3 –.**
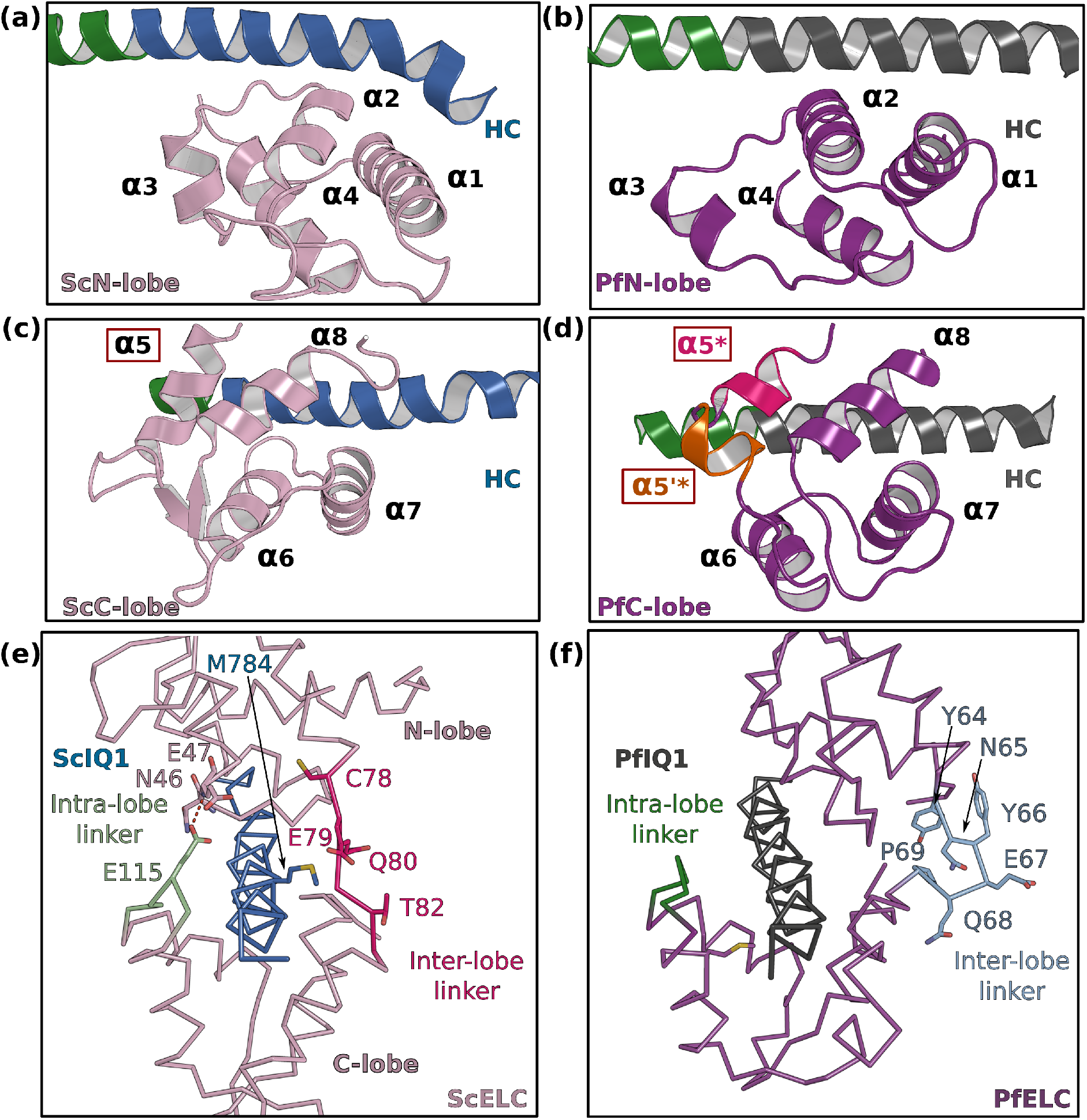
PfELC adopts an atypical fold. **(a)** The N-lobe of scallop myosin 2 ELC (ScMyo2) is compared to **(b)** the N-lobe of PfELC. The two lobes are conserved in structure, and adopt a closed conformation. In contrast, the C-lobe of ScMyo2 ELC **(c)** and PfELC **(d)** differ. The most divergent feature is the much shorter α5* helix comprised of only one turn and the presence of the short α5’* helix in PfMyoA. **(e)** shows the elongated interlobe linker of ScELC (red) compared with **(f)** the kinked, hairpin-like inter-lobe linker of PfELC (light blue). The specific structure of the inter-lobe linker makes the PfELC structure more compact than canonical ELCs and no direct interaction can occur between the lobes in PfMyoA to encircle the heavy chain, contrary to what is seen in the case of ScMyo2.

The sequence of the first IQ motif is adapted to the atypical features of PfELC. PfIQ1 is highly degenerate, containing none of the consensus IQ motif residues (IQxxxRGxxxR) (**Fig. 4a**; **Supplementary Fig. 4a, 4b, 4c**). In particular, PfIQ1 lacks the glutamine and the proximal arginine that make specific polar interactions with the C-lobe intra-lobe linker (between α6 and α7) in other myosins^20^ (**Fig. 4b**). In PfMyoA, these two residues are replaced by a valine (V781) and a glutamate (E785), respectively (**Fig. 4c**). While they also bind the intra-lobe linker of the semi-open C-lobe, the change in the nature of these interactions (**Fig. 4b, 4c**) results in a shift of the position of the intra-lobe linker relative to the HC helix (**Fig. 3e, 3f**). Interestingly, compared to other myosins, the IQ1 motif starts one residue earlier in the PfMyoA HC sequence because a HC residue is missing in the pliant region at the end of the Converter. This feature, in addition to a kink in the pliant region itself, contributes to a different orientation of the PfIQ1 HC helix (and thus the PfELC C-lobe) compared with other myosins when the Converter regions are superimposed (**Supplementary Fig. 7**). A different Converter/ELC interface is thus established (**Supplementary Fig. 7b, 7c**) and specific recognition of the semi-open C-lobe occurs both by its interaction with the Converter and with the HC helix. An atypical Trp (W777) in this helix strengthens the hydrophobic interactions between PfIQ1 and PfELC (**Fig. 4d, 4e**). Conventional myosins have a smaller side chain (I782) that interacts weakly with the C-lobe (**Fig. 4d**). In PfMyoA, W777 is sandwiched between the α5*, α5’* and α6 helices, thus interacting with the hydrophobic core of the PfELC C-lobe (**Fig. 4e**). These interactions between W777 and the α5* and α5’* helices are not present in the folded-back PR structure that displays a much larger kink at the pliant region and thus moves W777 away. The absence of electron density for the two helical turns of α5* and α5’* in this crystal structure indicates that lack of stabilization by the Converter and the W777 side chain affects the fold of these atypical PfELC structural elements (**Supplementary Fig. 2c, 6d, Fig. 5a**). These results demonstrate how the degenerate PfIQ1 motif is adapted to the peculiar structural features of PfELC, promoting specific recognition.

**Figure 4 –.**
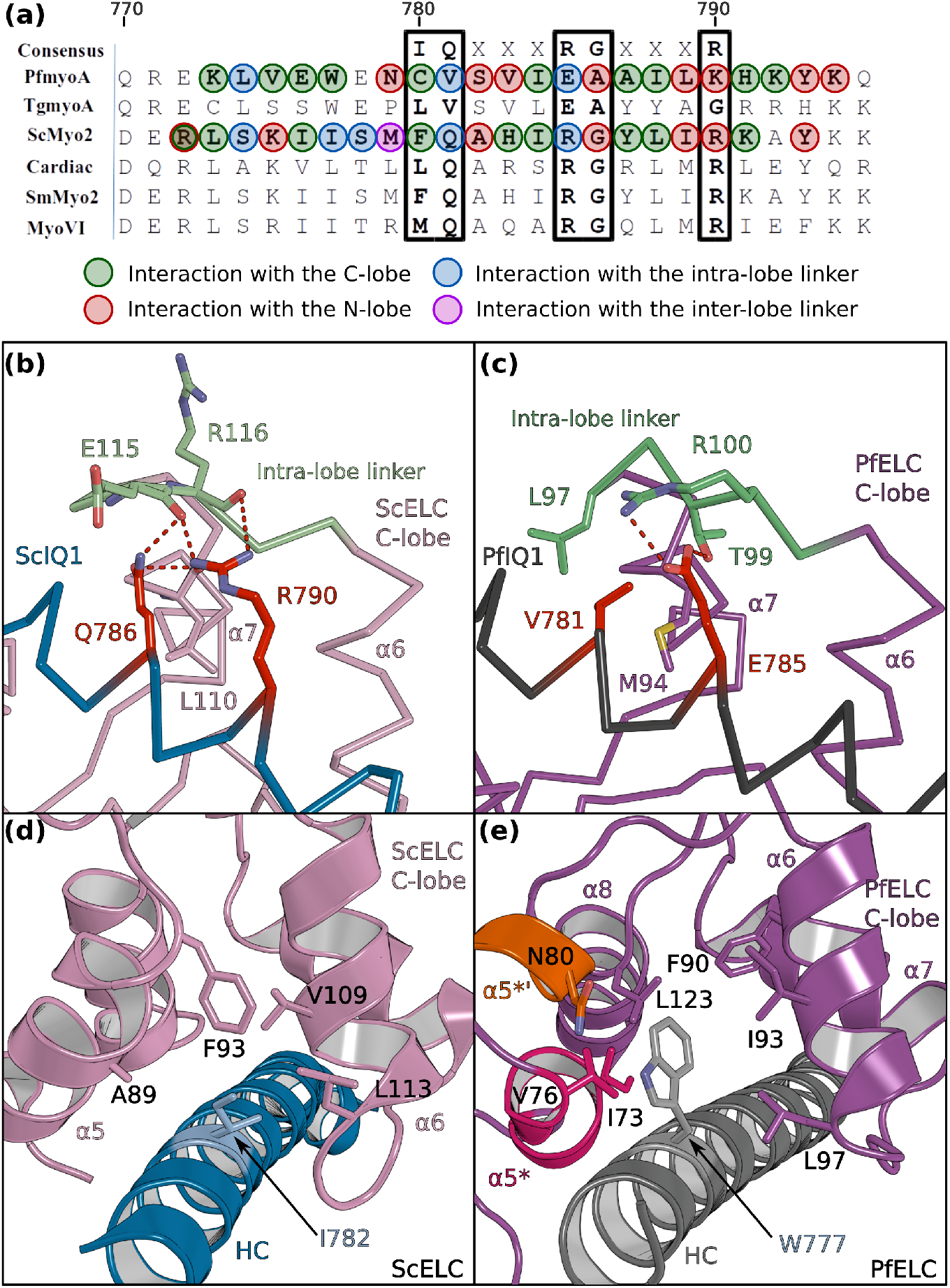
PfELC binds a degenerate IQ motif. **(a)** Sequence alignment of PfMyoA IQ1 to the consensus IQ motif and IQ motifs from other myosins: MyoA from T. gondii (TgmyoA); bay scallop (Argopecten irradians) myosin 2 (ScMyo2); human (Homo sapiens) β-cardiac myosin 2 (Cardiac); human Smooth muscle myosin 2 (SmMyo2); human myosin 6 (MyoVI). Consensus residues are contoured by a black box. **(b)** The intra-lobe linker of the scallop C-lobe interacts with the HC consensus residues with polar contacts. **(c)** In contrast, the intra-lobe linker of the Pf C-lobe is predominantly bound to the HC with apolar contacts. **(d,e)** Intra-lobe interactions maintain the semi-open C-lobe. Specificity in the recognition of the PfELC occurs via the W777 residue in PfMyoA. **(d)** In conventional myosins such as ScMyo2, a small side chain (I782) is found at the equivalent position, contributing to few interactions within the semi-open lobe. **(e)** In PfMyoA the bulky Trp (W777) is sandwiched between the α5*, α5’* and α6 helices, increasing the hydrophobic interactions with the PfELC C-lobe.

**Figure 5 –.**
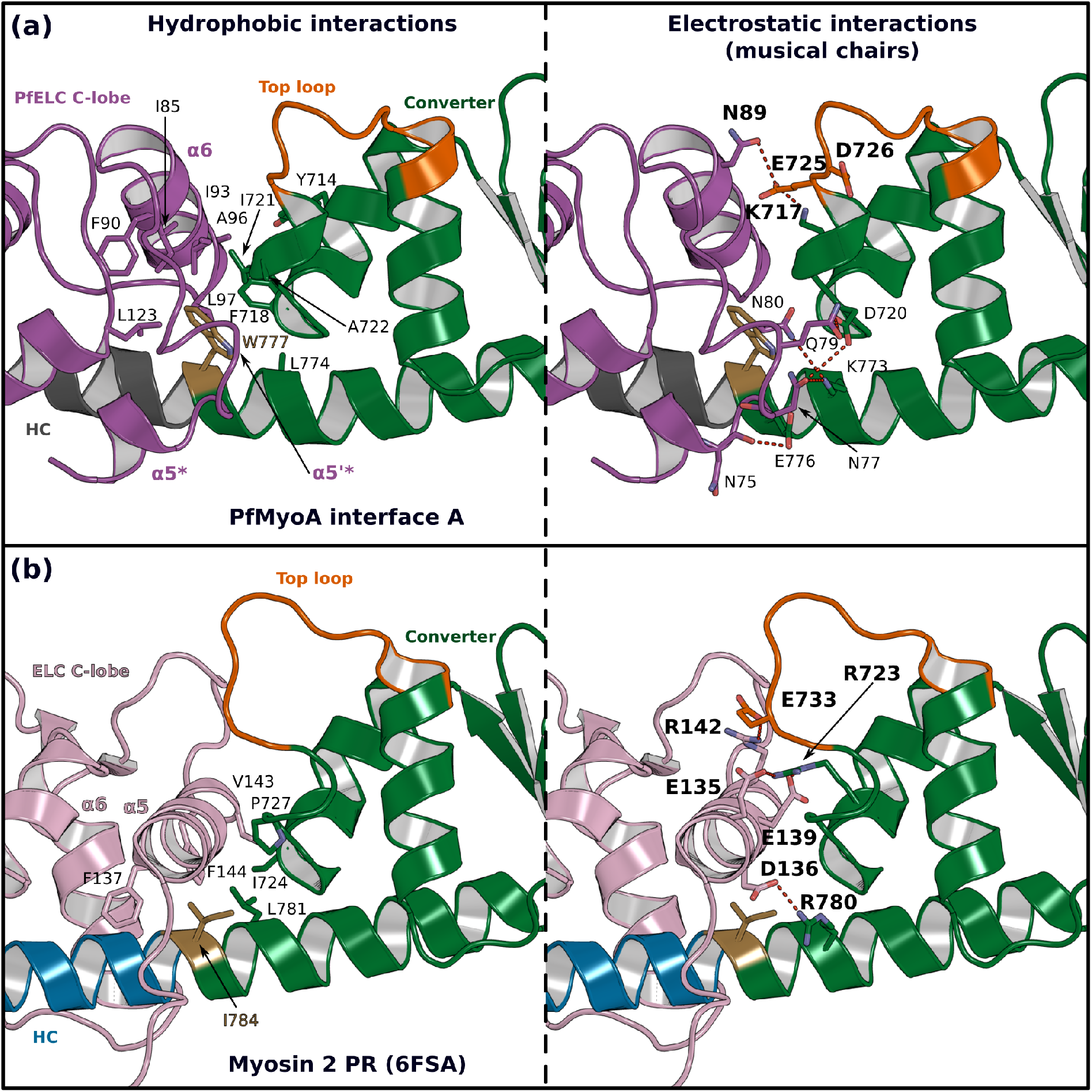
The atypical PfELC/Converter interface. Comparison of the Converter/ELC interface of **(a)** PfMyoA and **(b)** β-cardiac myosin. **(Left)**, hydrophobic contacts are displayed. **(Right)**, polar residues responsible for the contacts identified as “musical chairs” in molecular dynamics experiments^23^ are labeled.

### Compliance and hinge flexibility in the PfMyoA Lever Arm

The so-called pliant region between the Converter and the lever arm^14^ and the interfaces between light chains are hinges promoting lever arm flexibility^21,22^. The numerous differing structural features of the PfMyoA lever arm described above change these hinges of flexibility. We describe below the Converter/PfELC interface (interface A) seen in the PfMyoA•FL-PPS crystal structure, which is similar to the solution structure (**Supplementary Data 1**).

The shift in the PfELC position as compared with other myosins (**Supplementary Fig. 7**) results in a different Converter/ELC interface (**Fig. 5a, 5b**). In canonical myosins, the Converter/ELC interface is established between residues from the Converter and the α5 helix and includes mainly polar and few apolar interactions (**Fig. 5b**). In PfMyoA, a more rigid interface forms because it involves mainly hydrophobic contributions from the α5*, α5’* and α6 helices whose conformations are stabilized by W777 (**Fig. 5a**). The PfELC/MTIP interface is also mainly hydrophobic, involving residues from the MTIP α5 and α6 helices (H150, F151, I167, W171) and residues from the PfELC α1 helix and the following loop1 (**Supplementary Fig. 5f**). Interestingly, this interaction is only possible after the formation of a ^~^130° kink in the HC helix between the two IQs motifs of the PfMyoA lever arm. The kink is facilitated by proline (^HC^P802) and is similar in all molecules of the asymmetric units of the structural states we crystallized (**Supplementary Fig. 5g**). The contacts at the MTIP/PfELC interface contribute to stabilizing the kink and providing a specific conserved feature for this rather rigid lever arm.

To further investigate the dynamics of the PfMyoA lever arm, we performed molecular dynamics with explicit solvent on the PfMyoA•FL-PPS structure, an approach successfully used to investigate other myosins^22,9^. The first insight from this experiment is that priming of the PfMyoA lever arm is maintained throughout the time course. Even if the lever arm position slightly diverges at the beginning of the simulation, the priming is restored quickly and thus stays stable during the entire simulation (**Supplementary Movie 1**). As expected, the Converter/PfELC and PfELC/MTIP interfaces are rigid. All the hydrophobic interactions between PfELC and MTIP are maintained during the simulation and the Converter/PfELC interface is more rigid as compared with conventional myosins (**Supplementary Movie 1, 2**). In β-cardiac myosin, the Converter/ELC interface displays a controlled dynamic driven by labile polar interactions, the so-called “musical chairs”^22^. These interactions between charged residues are located at the pliant region, and involve the α5 helix of the ELC and the specific Top-loop of the Converter (**Fig. 5a, 5b, Supplementary Movie 2**). The dynamic associations between the musical chairs govern the flexibility of the Converter/ELC interface and the conformations allowed for the Top-loop^22^. In contrast, in PfMyoA, most of the Converter/ELC interface is hydrophobic and the bulky W777 introduces rigidity at this interface. The only musical chairs of the PfMyoA lever arm are located in two regions. Charged residues from the Top-loop (E725 and D726) interact alternatively with residues of the α6 helix (^PfELC^D88, ^PfELC^N89), restricting the Top-loop conformation. Other musical chairs involve residues from the Converter (D720; K773 and E776) with polar residues from the α5* and α5’* helices (^α5*^N75; ^α5’*^N77; ^α5’*^E78; ^α5’*^Q79) (**Fig. 5a; Supplementary Movie 2**). We conclude that the priming seen in the PfMyoA•FL-PPS crystal structure is stable, and that the PfMyoA lever arm is more rigid than in conventional myosins.

### Motor domain /Lever arm interactions stabilize the atypical PfMyoA priming

The atypical priming of the PfMyoA lever arm in the PPS state results from several interactions between the lever arm and the motor domain that are not present in conventional myosins. Conformational changes at the end of the SH1 helix are required to allow an additional Converter rotation so that it can reach the PfMyoA motor domain surface with which it interacts (**Supplementary Fig. 4d**). These interactions involve two interfaces: interactions of PfELC with Loop-1 and the β-bulge and a specific interface between the Converter and the N-terminal subdomain (N-term) (**Fig. 6a, 6b**). Interestingly, interactions between PfELC, Loop-1 and the β-bulge are highly labile during molecular dynamics (**Supplementary Movie 1**), but the interface between the Converter and the N-term subdomain is highly stable and maintained throughout the time course (**Supplementary Movie 1**).

**Figure 6 –.**
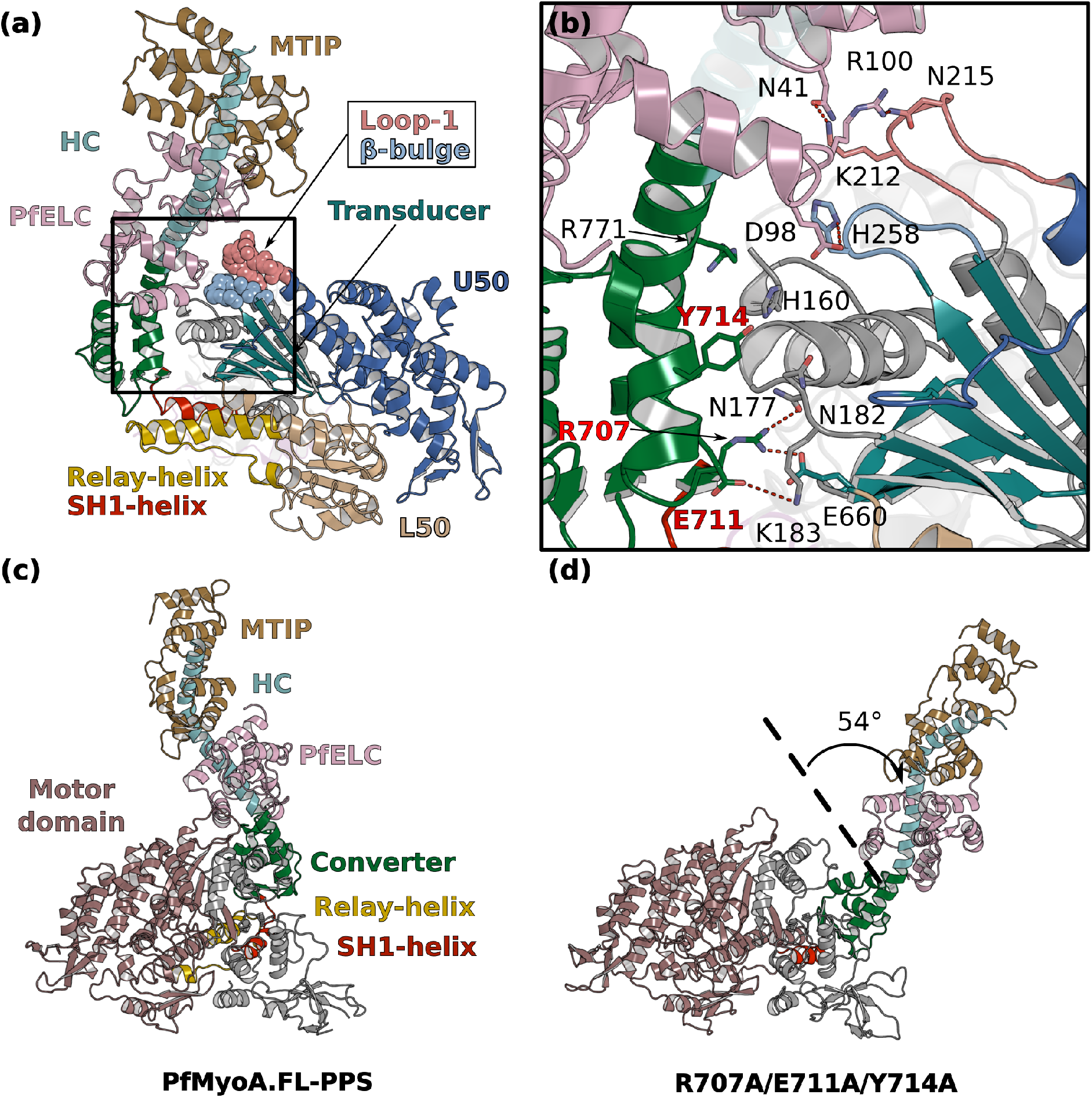
Interactions stabilizing the priming of the PfMyoA PPS state. **(a)** Overall structure of PfMyoA•FL-PPS. The priming of the lever arm is stabilized by interactions between elements of the lever arm and the motor domain (boxed in black). **(b)** Zoom on the Lever arm/motor domain interface. The residues involved in the interaction are labeled. Key residues that have been mutated to disrupt the interface (see **Fig. 6d**) are labeled red. **(c,d)** MD simulations comparing the WT and triple mutant R707A/E711A/Y714A. **(c)** The primed PPS state is stable during the entire duration of the simulation (320 ns). **(d)** In contrast, the priming is lost with the triple R707A/E711A/Y714A mutant and the position of the lever arm deviates by up to 54°.

The Converter/N-term subdomain interface can be subdivided into two regions. The first region involves a set of two polar interactions involving ^Converter^R707/^Nterm^N177/^Transducer^E660 and ^converter^E7n11/^Nterm^K183 and also hydrophobic interactions established between ^Converter^Y714 and ^Nterm^V181 and ^Nterm^N182 (**Fig. 6b**). These interactions are maintained throughout the duration of the molecular dynamics (**Supplementary Movie 1**). While the position of ^Converter^Y714 shifts, it remains part of the interface. The second region consists in cation/π-stacking interactions between ^Converter^R771 from the last helix of the Converter and ^Nterm^H160. This interaction is also stable throughout the time course of the dynamics, while ^Converter^R771 occasionally establishes an interaction with ^Nterm^D159 (**Supplementary Movie 1**).

To test the hypothesis that this Converter/N-term interface is key for stabilization of the atypical priming of PfMyoA in the PPS state, we designed the triple mutant R707A/E711A/Y714A *in silico*. As expected, molecular dynamics show that the three mutations destabilize the Converter/Nterm subdomain interface and cause a loss of priming, demonstrating that this interface stabilizes the atypical priming of PfMyoA in the PPS state (**Fig. 6c, Supplementary Movie 3,4**).

The consequences of this atypical priming on the motor properties of PfMyoA were tested with two triple mutants: R707A/E711A/Y714A and R707L/E711R/Y714A (priming AAA and LRA mutants). The two sets of mutations are expected to disrupt the Converter/MD interface and thus to reduce the priming of the PPS. Both triple mutants greatly impact and tune the properties of the motor similarly: they reduce the maximum actin-activated ATPase to ^~^30% of WT, and decrease both *in vitro* motility speed and the ADP release rate to a similar extent (^~^35% of WT) (**Fig. 2b-e, Supplementary Figure 8a-d, Supplementary Table 2**). The decreased ADP-release rate implies that these Converter residues play a role in the stability of the Strong-ADP state the motor populates when attached to actin. It is likely that a smaller step size also contributes to the reduced speed at which the mutant motor moves actin, because *in silico* studies show that the AAA mutant fails to maintain the WT lever arm orientation, although the Converter stays oriented in a primed orientation (**Figure 6c**).

### The PfELC is essential for plasmodium invasion

As described above, the PfELC is uniquely suited to bind to the degenerate IQ motif and is also engaged in stabilizing the unusual priming of this motor, as well as being an essential element to maintain the rigidity of the lever arm. To validate this essential role and to test whether alternate light chains might compensate for PfELC function (as exists in *Toxoplasma gondii^23^*), we generated a conditional knockout for the *Pfelc* gene in *Plasmodium falciparum*. Using a combined process of drug selection-linked integration and engineering of artificial Cre recombinase sites into the *PfELC* gene via synthetic introns^24,25^ (**Supplementary Figure 8e**), we were able to selectively induce PfELC truncation at the protein C terminus (lacking amino acids 93-134, predicted to render it non-functional). Integration of the introns did not effect parasite asexual growth (**Figure 7a**). Treatment with rapamycin, inducing diCre dimerization and gene excision, led to gene truncation (as validated by PCR (**Supplementary Figure 8f**). Truncation was also confirmed by immunoblot and immunofluorescence assay (**Figure 7b, 7c, 7d**). Analysis of rapamycin treated parasites was associated with a near complete ablation of red blood cell invasion over 60 hours of an asexual growth cycle, on a par with inhibition levels seen with the non-specific invasion inhibitor heparin^26^ (**Figure 8e**). Further analysis over a single 48 hours or double 96 hours cycle showed that this inhibition of invasion was near total (**Figure 7f, 7g**) suggesting residual invasion after a single cycle is likely the result of incomplete gene excision. The level of invasion retardation following loss of PfELC suggests that, like the heavy chain PfMyoA^9^, PfELC is essential for asexual blood-stage life cycle progression. Unlike *T. gondii*, there is no evidence for redundancy in essential light chains. Both PfMyoA motor domain and PfELC thus represent attractive targets for arresting parasite invasion in pathogenic blood stages.

**Figure 7 –.**
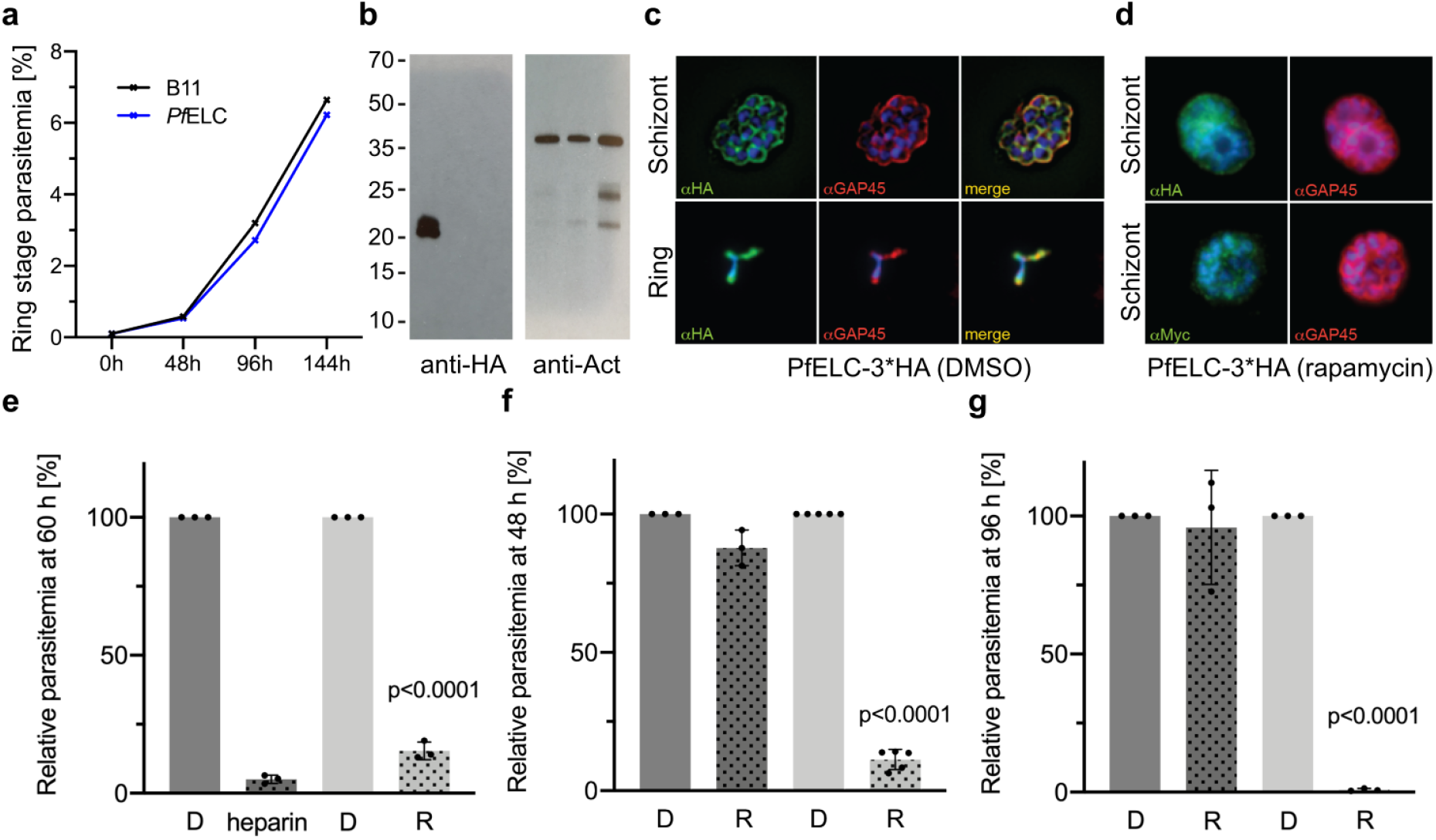
PfELC is essential for parasite invasion of the red blood cell. **(a)** Growth curve comparing B11 wildtype and PfELC-3xHA parasite over the course of 3 cycles indicates no detrimental effect of genetic modification in the pfelc locus. Mean ring-stage parasitemia is shown of two biological replicates as determined by FACS. **(b)** Western blot analysis of parasite extracts separated by SDS-PAGE probed with anti-HA and anti-actin antibodies as a loading control. PfELC-3xHA runs at the expected MW of ^~^ 21 kDa in mock DMSO treated samples (D) and the HA signal is lost upon treatment with rapamycin (R), indicating successful excision. **(c-d)** Representative IFAs of schizont and ring-stage parasites co-labelled with the IMC marker GAP45. Peripheral staining is lost after treatment with rapamycin and a diffuse/punctuate pattern is seen with anti-cMyc antibodies. **(e)** R-treated PfELC 3xHA parasites are significantly impaired in invading red blood cells as determined by FACS analysis of ring-stage parasites over 60 hours comparable to that seen with the generic inhibitor heparin. A small amount of residual invasion seen after 48h **(f)** disappears after 96h post treatment **(g)** suggesting absence of PfELC results in complete ablation of invasion. D-treated parasites show no invasion defect.

## Discussion

The full-length PfMyoA structures reveal how the atypical lever arm of this myosin is part of the adaptations tuning its motor cycle. Unexpectedly, the lever arm of PfMyoA is ^~^30° more primed upon reattachment to the actin track than it is in conventional class II myosins. Designed from structural insights, *in vitro* and *in silico* experiments reveal that specific interactions stabilize the priming of PfMyoA and define specific adaptations tuning the properties of this motor. These interactions are not present in Scallop Myosin 2 (**Supplementary Figure 9a, 9b**). Myo10 and Myo1c are two motors for which the lever arm priming position is also large in the pre-powerstroke state^16,27^, but their lever arm orientation is stabilized by a different set of interactions when compared with PfMyoA (**Supplementary Figure 9a, 9c, 9d**). The contribution of the Transducer in the stabilization of the primed lever arm also differs in the three myosins. It is thus tempting to speculate that the nature of these interactions will tune each motor for its specific functions. Increasing the amplitude of the lever arm priming could be a way to lengthen the step size of the motor and thus to increase the speed at which it can move actin. This is consistent with the reduced speed of *in vitro* motility obtained with PfMyoA mutants designed to reduce the priming of the lever arm as well as data for Myo10 indicating that this motor is tuned for performing large steps on actin bundles^16^. Specific interactions with the Transducer may also be involved in enlarging the stroke and tuning the orientation of the powerstroke for high duty-ratio motors able to resist rearwards load, as with the Myo1b motor that anchors the cell membrane to the cytoskeleton^28^. As it is the case for class I Myosins^29,30^, the duty-ratio of PfMyoA is regulated by a specific N-terminal HC extension^9^. The observation that Converter mutations involved in priming PfMyoA also decrease the ADP-release rate suggests that these Converter residues might be involved in facilitating the transitions occurring during the PfMyoA powerstroke.

Atomic structures of the PfMyoA lever arm indicate how degenerate IQ motifs of the HC are recognized by PfELC and MTIP, as well as how unusual apolar interactions occur between PfELC and the Converter. These unusual features could not have been predicted from recently solved TgMyoA structures of the lever arm bound to TgELC or TgELC2^31^, because TgELC is a poor model for PfELC considering the low sequence identity (20 and 21% between PfELC and TgELC and TgELC2, respectively). Interestingly, TgELC1 and TgELC2 are homologous to classical ELCs and do not display the specific inter-lobe linker nor the atypical α5* and α5’* helices found in PfELC, which are also part of the interface with the Converter. This structural difference illustrates again that Myosin A from different organisms differ in detail due to the high divergence of the different taxa from the Apicomplexa phylum. The structure of PfELC and its specific association with the HC and MTIP also explains the cooperative binding of these elements and why PfELC does not bind to the HC in the absence of MTIP^5^. Indeed, the structures reveal a sequence shift near the pliant region resulting in a specific orientation of the HC helix in order to stabilize the Converter/PfELC interface (**Supplementary Figure 7**). The PfELC α5* and α5’* helices are involved in this interface but may not be properly folded prior to its establishment. In addition, PfELC cannot surround the IQ1 motif due to its unusual interlobe linker, which may contribute to diminishing a stable association with the HC. According to this hypothesis, PfMTIP may need to bind first to the IQ2 motif and its presence could add stabilizing interactions to cooperatively recruit PfELC to the PfIQ1 motif via formation of the interfaces with PfMTIP and the Converter, allowing stabilization of the PfELC α5* and α5’* helices. PfELC is thus a weak point in the assembly of the fully functional motor complex, and compounds targeting PfELC may be a good strategy for decreasing motor activity. PfMyoA lacking the ELC moves actin at half the speed of the motor containing both light chains^5^. Moreover, studies with skeletal muscle myosin showed that removal of its essential light chain reduced isometric force by over 50%^32^. Either or both of these impairments likely contribute to the finding that the PfELC is essential for red blood cell invasion.

Taken together, the data presented here reveal important new findings about the PfMyoA motor, and how the lever arm is involved in tuning the specific mechano-properties of this atypical myosin. While the N-terminal HC extension of PfMyoA is involved in regulating the transition between the Strong ADP and the Rigor states, the sequence adaptations located in the Relay, the SH1-helix or the Wedge^9^ influence other structural transitions essential for motor function. We show that the overall stabilization of the PPS differs in this myosin compared with all other myosins in the superfamily. Control of the recovery and powerstroke transitions differ in detail although the overall sequence allows conservation of the essential properties of a motor: the ability to couple the lever arm priming with ATP hydrolysis and control of the powerstroke via interaction with F-actin, associated with sequential controlled release of the products phosphate and ADP.

Lastly, we have also demonstrated that PfELC, like PfMyoA^9^, is essential in parasite asexual invasion of the red blood cell and therefore a second attractive target within the glideosome for targeting malaria parasite asexual replication, where absence of either alone or in combination would block red blood cell entry. While strategies to disrupt the glideosome have already been investigated^33^, inhibition of the PfMyoA motor^9^ or the association of the PfELC light chain opens up new strategies towards therapeutic solutions. Importantly, the structures described here provide a precise blueprint for designing just such small molecules that could prevent binding of PfELC or target PfMyoA full length motor activity and thus diminish glideosome function. These results inform the design of new therapies that specifically target lifecycle progression in the pathogenic blood stages of *Plasmodium*.

## Methods

### Expression constructs

Full-length PfMyoA heavy chain (PlasmoDB ID PF3D7_1342600/ GenBank accession number XM_001350111.1), with Sf9 cell preferred codons, was cloned into the baculovirus transfer vector pFastBac (pFB) (Thermo Fisher). A 13 amino acid linker separates the C-terminus of the PfMyoA heavy chain from an 88 amino acid segment of the Escherichia coli biotin carboxyl carrier protein {Cronan, 1990 #1}, which gets biotinylated during expression in Sf9 cells, followed by a C-terminal FLAG tag for purification via affinity chromatography. Heavy chain mutants (point mutants E6R, T586F, S691G; triple mutants R707A/E711A/Y714A and R707L/E711R/Y714A) were generated on this backbone using site directed mutagenesis. An N-terminal truncation of MTIP (MTIP-Δn, residues 1-E60 deleted and an N-terminal HIS tag), and an N-terminal truncation of the heavy chain (ΔN-terminal HC extension (ΔNter), residues 1-S20 deleted) were also cloned for use in constructs that were crystallized. Recombinant baculovirus was produced using the Bac-to-Bac Baculovirus expression system (Thermo Fisher). The mouse utrophin (NP_035812) clone was a gift from Kathleen Ruppel and James Spudich. It was modified so that utrophin residues 1-H416 were followed by C-terminal biotin and FLAG tags. It was cloned into pFastbac for production of recombinant baculovirus and subsequent expression in Sf9 cells.

### Myosin expression and purification

For biochemical characterization, full-length PfMyoA heavy chain mutant constructs were co-expressed with the chaperone PUNC and the light chains (PfMTIP and PfELC) in Sf9 cells as described in^5^. Two constructs were expressed for crystallization. In one, the WT heavy chain, PfELC, and MTIP-Δn were co-expressed with the chaperone PUNC. In the second, ΔNter heavy chain, PfELC, and MTIP-Δn were co-expressed with the chaperone PUNC. The cells were grown for 72 h in medium containing 0.2 mg/ml biotin, harvested and lysed by sonication in 10mM imidazole, pH 7.4, 0.2M NaCl, 1mM EGTA, 5 mM MgCl2, 7% (w/v) sucrose, 2 mM DTT, 0.5mM 4-(2-aminoethyl)benzenesuflonyl fluoride, 5 μg/ml leupeptin, 2mM MgATP. An additional 2 mM MgATP was added prior to a clarifying spin at 200,000×g for 40 min. The supernatant was purified using FLAG-affinity chromatography (Sigma). The column was washed with 10mM imidazole pH 7.4, 0.2M NaCl, and 1 mM EGTA and the myosin eluted from the column using the same buffer plus 0.1 mg/ml FLAG peptide. The fractions containing myosin were pooled and concentrated using an Amicon centrifugal filter device (Millipore), and dialyzed overnight against 10mM imidazole, pH 7.4, 0.2M NaCl, 1mM EGTA, 55% (v/v) glycerol, 1 mM DTT, and 1 μg/ml leupeptin and stored at −20 °C. Utrophin purification was essentially the same as for myosin but without the MgATP steps. Skeletal muscle actin was purified from chicken skeletal muscle tissue essentially as described in^34^.

### Biochemical assays

Unloaded and loaded *in vitro* motility assays, actin-activated ATPase assays, and transient kinetic measurements were performed as described in^9^.

### Crystallization and data processing

Crystals of PfMyoA•FL-PR (10 mg.ml^−1^) (Type A) were obtained at 4°C by the hanging drop vapor diffusion method from a 1:1 (v:v) of protein with 2 mM MgADP and precipitant containing 1.9M Ammonium sulfate, 0.1M Sodium HEPES pH 6.8, 2% PEG400. Crystals of PfMyoA•FL-PPS (Type B) were obtained at 17°C by the sitting drop vapor diffusion method from a 1:1 mixture of protein (10 mg.ml^−1^) with 2 mM MgADP.VO4 and precipitant containing 1M Na Citrate tribasic; 0.01M Na Borate pH 8.0. Crystals of PfMyoA•ΔNter-PR were obtained at 4°C by the hanging drop vapor diffusion method from a 1:1 mixture of protein (10 mg.ml^−1^) with 2 mM MgADP and precipitant containing 2.0M Ammonium sulfate, 0.1M Sodium HEPES pH 7.5, 6% PEG400.

Crystals were transferred in the mother liquor containing 30% glycerol before and flash freezing in liquid nitrogen. X-ray diffraction data were collected at the SOLEIL synchrotron, on PX1 beamline (λ = 0.906019 Å for type A, λ = 0.978570 Å for type B, λ = 0.978570 Å for type C), at 100 K. Diffraction data were processed using the XDS package^35^ and AutoPROC^36^. Crystals type A and C belong to the P2_1_2_1_2_1_ space group, crystals type B belong to the P2_1_2_1_2 space group, with one molecule per asymmetric unit for type A and C and two molecules per asymmetric unit for type B. The data collection and refinement statistics for these crystals are presented in (**Supplementary Table 1**).

### Structure determination and refinement

Molecular replacement was performed with the PfMyoA motor domain coordinates (21-768) (PDB code 6I7E for the PPS; PDB code 6I7D chain A for the PR^9^) with Phaser^37^. The structure in the PR state (PDB code 6I7D, chain A) without ligand and water was used as a target model for type A and C crystals. PfMyoA motor domain in the PPS state (PDB code 6I7E) was used as a target model for type B crystals. Manual model building was achieved using Coot^38^, the structure of MTIP complexed with the cognate IQ motif peptide (PDB code 4OAM^13^ was used to rebuild MTIP. Refinement was performed using Buster^39^. The statistics for most favored, allowed and outlier in Ramachandran angles are for each crystal type respectively (in %): 96.40, 3.41, 0.19 for PfMyoA•FL-PR; 91.93, 6.80 and 1.27 for PfMyoA•FL-PPS; 93.12, 5.92, 0.96 for PfMyoA•ΔNter-PR.

### SAXS experiments

SAXS data were collected at the SOLEIL synchrotron, on the SWING beamline (λ = 1.03319947498 Å). Purified PfMyoA/ELC/MTIP-Δn was extensively dialyzed against 10mM Hepes pH 7.4, 100 mM NaCl, 1 mM DTT, 1 mM NaN3 (without any ATP) in order to remove nucleotide. We prepared two samples PfMyoA-PR and PfMyoA-PPS. Both were subsequently incubated with 2 mM MgADP for 20 min on ice, and then we added 2 mM vanadate for PfMyoA-PPS, but not for PfMyoA-PR. All samples were centrifuged at 20,000×g for 10 min at 4 °C prior to the analysis. 40 μl of the protein at 2, 5 and 9 mg.ml^−1^ (17, 41 and 75 μM, respectively) were injected between two air bubbles using the auto-sampler robot. Thirty-five frames of 1.5 s exposure were averaged and buffer scattering was subtracted from the sample data. As all 2, 5 and 9 mg ml^−1^ curves displayed no traces of aggregation, only the 9 mg.ml^−1^ curve was used for further analysis because of the higher signal/noise ratio. The theoretical SAXS curves were calculated with CRYSOL and compared based on the quality of their fits against the different experimental curves. We computed the SAXS envelopes of the two samples with GASBOR, two programs from the ATSAS suite^40^. The analysis and the fit of the computed SAXS envelopes and the X-ray structures were performed with FoXS^41^. Furthermore, to analyze if we have a mixed population of PR-Closed and PR-open conformations of PfMyoA-PR in solution we used Oligomer^40^. The model of PR-open conformation used in this study was generated by Swiss model^42^ after manual positioning of the lever arm, as found in the PPS structure in which the pliant region is not kinked.

### Molecular dynamics

Molecular dynamics experiments were performed with a procedure close to^22^. All the systems (WT + mutants) were built with the CHARM-GUI ^43,44^, with the Solution Builder module. The entire proteins were relaxed in a box containing explicit water (TIP3P) and salt (150 mM KCl) at 310.15 K in the CHARMM36m force field^45^. The duration of the simulations was 320 ns in GROMACS (version 2018.3)^46^.

### Generation of the PfELCloxP construct

A gene fragment (GeneArt) was synthesized comprising a 454 bp targeting sequence of the *pfelc* gene (3D7_1017500), a loxPint module replacing the second intron and the last 126 bp (codon-optimized) exon, encoding the C-terminal end of PfELC (Ile93 to Ile134) and a triple hemagglutinin tag. The fragment was cloned into the NotI/AvrII site of a modified version of the conditional KO vector^47^, containing the SLI elements^25^ and a cMyc/Flag tag. The resulting plasmid was purified from *E. coli* using the Qiagen Plasmid Maxi kit.

### *P. falciparum* culture and transfection

*P. falciparum* B11 parasites expressing the DiCre recombinase^33^ were cultured in RPMI 1640 medium containing 0.5% w/v AlbumaxII and at 4% haematocrit using human erythrocytes (blood group 0^+^) according to standard procedures^48^. Ring-stage parasites were synchronised with 5% sorbitol (w/v) and transfected with 100 μg of purified pARL PfELCloxP plasmid. Transgenic parasites harbouring the episomal plasmid were initially selected with 2.5 nM WR99210 (Jacobus Pharmaceuticals) for 7 days and integration into the genomic locus was achieved by G418 treatment (400 μg/ml) for 10 days.

DiCre-mediated excision was achieved by treating synchronised ring-stage parasites with 100 nM rapamycin (Sigma) in DMSO for 14-16 h. Additionally, parasites were mock-treated with 1% (v/v) DMSO as a negative control. Cultures were washed three times with culture media to remove rapamycin and DMSO, respectively. Treated parasites were transferred into 96-well plates at 0.1% parasitemia (in triplicate) containing fresh red blood cells at 0.1% hematocrit and were allowed to proceed to the next ring-stage cycle for FACS analysis. Samples for gDNA extraction and PCR amplification, Western blot and immunofluorescence analysis were taken towards the end of the treatment cycle or subsequent cycles. Parallel assays were carried out with a negative control, B11 parasites treated with heparin throughout (Pfizer, 1:25). For these assays, parasites were treated with rapamycin and washed as above, then transferred to 48-well plates at 1% parasitaemia (in triplicate) containing fresh red blood cells at 5% hematocrit and parasitaemia was analysed as above, at around 60 h post-treatment.

### Genotyping, Western blot analysis and immunofluorescence assays

DNA was extracted using the PureLink Genomic DNA Mini kit (Invitrogen) for genotyping by PCR using primers P1 5’ CATTACTTTAATTTTTATACTACTGTTTATTTTTACAGTAC 3’, P2 5’ CTAATCCTATTATTT AAATATTTCATATTTTTTAAACATAGATGG 3’, P3 5’ GGCCAGCCACGATAGCCGCGCTGCCTCG 3’, P4 5’ CTTGTCGTCATCGTCTTTGTAGTCCTTGTC 3’ and P5 5’ CAGGAAACA GCTATGACCATG 3’ and KOD Hot Start DNA Polymerase (Millipore).

Schizonts were lysed with 0.2% saponin/PBS, washed three times with PBS (supplemented with cOmplete™ EDTA-free protease inhibitors, Roche) and resuspended in 1x SDS loading dye. Proteins were separated on NuPAGE™ Novex^®^ 4-12% Bis-Tris protein gels in MES buffer (Life Technologies) and were transferred onto nitrocellulose membranes using the iBlot^®^ system (Life Technologies). Membranes were blocked in 5% skim milk/PBS (w/v) and probed with the following antibodies diluted in 5% skim milk/PBS (w/v): anti-HA (1:4000, clone C29F4, Cell Signaling), anti-cMyc (1:100, clone 9E10, Invitrogen) and rabbit anti F-actin ^49^(1:1000), Horseradish peroxidase-conjugated goat anti-rabbit and goat anti-mouse antibodies were used as secondary antibodies and diluted 1:10,000 (Jackson IR).

Late stage parasites were treated with cysteine protease inhibitor E64 (10 μM) for 4 h prior to fixation with 4% PFA/0.0025% glutaraldehyde/PBS for 1 h. Fixed cells were permeabilised with 0.1% TX-100/PBS for 15 min, blocked in 3% BSA and probed with anti-HA (clone 12CA5, Roche), anti-cMyc (clone 9E10, Invitrogen) and anti-GAP45^50^ antibodies at 1:500 in 3% BSA/PBS for 1 h, followed by three washes in PBS. Secondary Alexa Fluor^®^ conjugated antibodies (Invitrogen) and DAPI (4’,6-diamidino-2-phenylindole) were diluted at 1:4000 and incubated for 1 h, followed by three washes in PBS. Images were acquired with an OrcaFlash4.0 CMOS camera using a Nikon Ti Microscope (Nikon Plan Apo 100 × 1.4-N.A. oil) and Z-stacks were deconvolved using the EpiDEMIC plugin with 80 iterations in Icy (DeChaumont et al, 2012). Images were processed in Fiji/Image J^51^ Representatives of 3-5 biological replicates are shown. Significance assessed by unpaired t-test, two tailed.

### Growth analysis by FACS

Tightly synchronised ring-stage parasites were diluted to 0.1% parasitemia (in triplicate) for growth curves and transferred into 96-well plates containing fresh red blood cells at 0.1% haematocrit. Parasites were allowed to proceed to the next ring-stage cycle, were stained with SYBR Green (1:10,000, Sigma) for 10 min and washed three times with PBS prior to collection by flow cytometry using a BD LSRFortessa™. A 100,000 cells were counted and FCS vs SSC was used to gate for red blood cells, SSC-A vs SSC-W for singlets and FSC-H vs SYBR-A was applied to gate for infected red blood cells. Flow cytometry data was analysed in FlowJo. Biological replicates are indicated in the figures.

## Data availability

The atomic models are available in the PDB, www.pdb.org, under accession numbers PDB 6YCX, 6YCY and 6YCZ for the PfMyoA•FL-PR, PfMyoA•FL-PPS and PfMyoA•ΔNter-PR, respectively.

## Acknowledgments

We are grateful to beamline scientists of PX1 (SOLEIL synchrotron) for excellent support during data collection. We thank Margaret A. Titus for critical reading of the manuscript. This work was supported by National Institutes of Health multi-PI grant AI 132378 to K.M.T and A.H and Human Frontier Science

Program grant to J.B. and A.H. (RGY0066/2016). Parasite work was funded through an Investigator Award to J.B. (100993/Z/13/Z) and PhD studentship to T.C.A.B. (109007/Z/15/A) both from Wellcome.

## Author contributions

D.M., J.P.R., D.A., J.R.P., K.M.T. and A.H. designed the research. E.B.K. expressed and purified protein for crystallization. D.M. crystallized PfMyoA constructs in the different states. D.M. solved the structures and performed refinement with the help of J.R.P. D.A. performed molecular dynamics. J.P.R. performed *in vitro* functional assays. S.H., T.C.A.B and J.B. designed and executed parasite work. D.M., J.P.R., D.A., J.R.P., K.M.T. and A.H. analyzed the results and wrote the paper.

