## Supplementary data for "Structure of Full Length Plasmodium Myosin A and its light chain PfELC, dual targets against malaria parasite pathogenesis"

### Supplementary Information

**Moussaoui *et al.*, 2020**

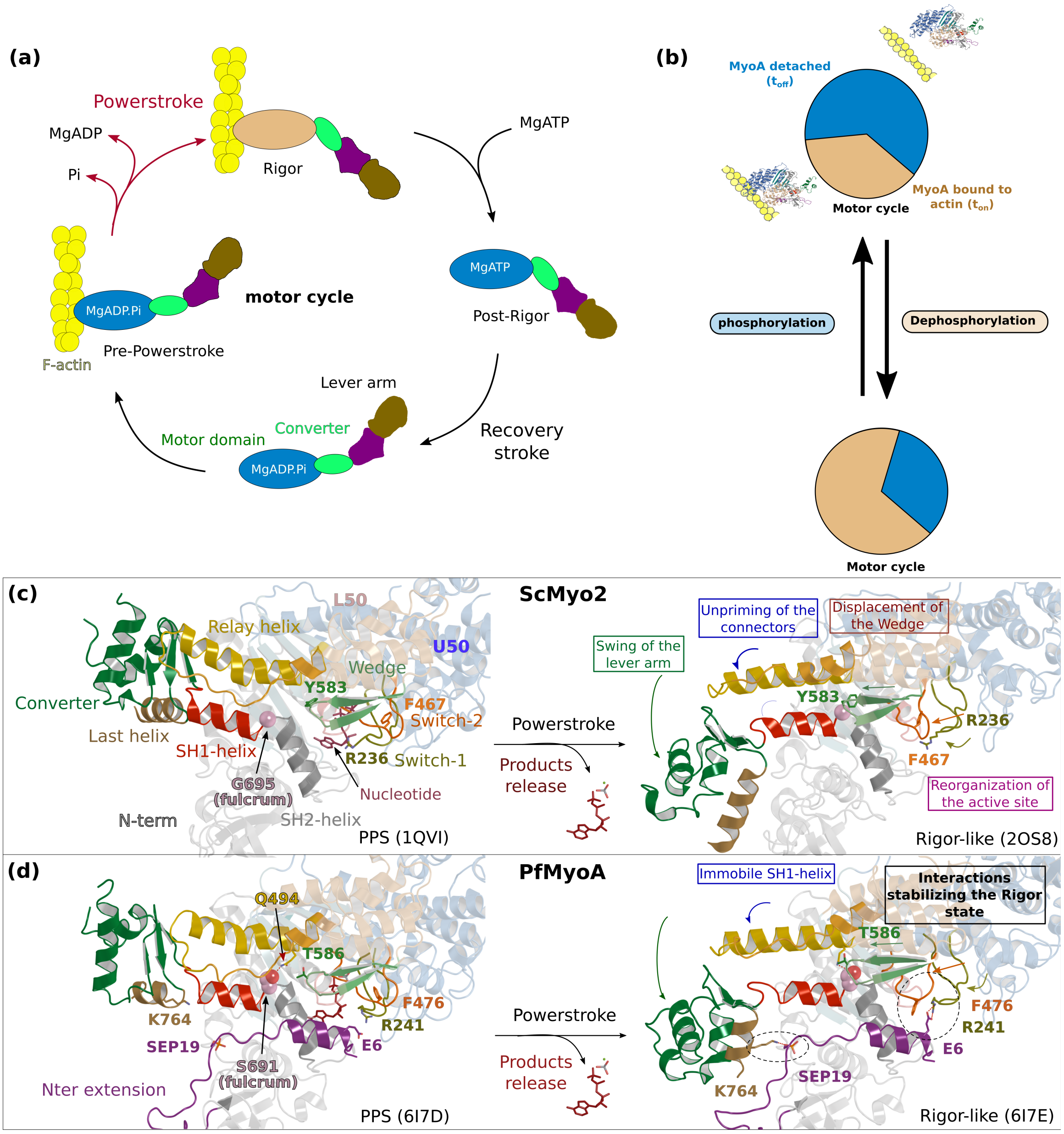

*<u>Supplementary Figure 1 – The atypical and tunable mechanical cycle of PfMyoA.</u>* ***(a)*** *Motor cycle of myosin motors. The nucleotide free state, strongly bound to actin, is called Rigor. Binding of ATP in the Rigor state detaches the head from actin. Upon detachment, the motor first populates the Post-Rigor state (PR) with ATP bound. Isomerization towards the Pre-Powerstroke state allows ATP hydrolysis. This state binds weakly to actin. The sequential release of the products of hydrolysis drives the lever arm swing and thus the powerstroke, which generates force. Detachment of the motor upon ATP binding starts a new cycle.* ***(b)*** *PfMyoA properties are tuned by a phosphorylation in the N-terminal extension of the motor domain (SEP19). When the motor is phosphorylated, it moves actin with a high velocity spending a short fraction of the cycle strongly bound to actin. When the motor is dephosphorylated, it produces more force at the expense of speed.* ***(c)*** *and* ***(d)*** *show the mechanism of force production in Scallop myosin 2 (ScMyo2) and PfMyoA respectively. In ScMyo2 the mobility of the SH1-helix is a key element for the mechanism driving the powerstroke of most myosins and requires the presence of a conserved glycine (G695), the so-called fulcrum, at the basis of the SH1-helix[1,2,3]. In PfMyoA, the SH1-helix is immobile, due to the presence of a Serine at the fulcrum (S691). The lack of mobility of the fulcrum is compensated by a phosphorylatable N-terminal extension (SEP19) and sequence adaptations in the connector.*

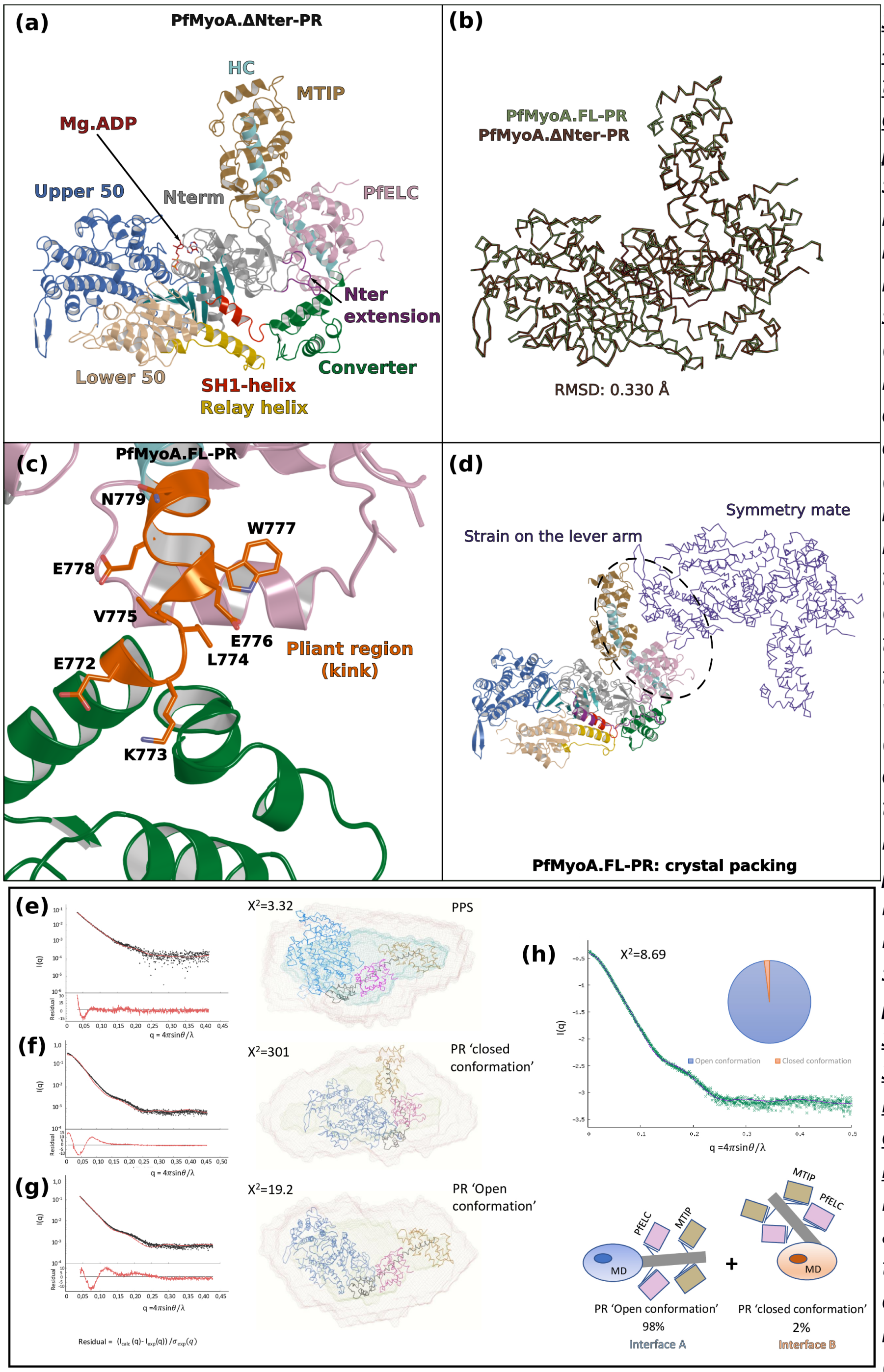

<u>*Supplementary Figure 2 – The crystal structure of the PfMyoA PR state displays a kink at the pliant region.*</u> ***(a)*** *Overall structure of PfMyoA•ΔNter-PR.* ***(b)*** *PfMyoA•ΔNter-PR and PfMyoA•FL-PR superimpose perfectly (rmsd 0.330 Å), indicating that the deletion does not change or alter the protein fold.* ***(c)*** *The PR state of PfMyoA•FL displays a kink in the lever arm at the pliant region (orange). The Converter thus becomes far from the α5* and α5*' helices which are destabilized (not modeled since no density is indicated in the electron density map).* ***(d)*** *Crystal packing shows that the kinked lever arm is involved in a large surface of the crystal packing.* ***(e), (f), (g)*** <u>*Small-Angle X-ray Scattering experiment investigating the conformation of PfMyoA in solution.*</u> ***(e)*** *When the motor is bound to ADP and Pi analogs, the theoretical curve computed from the PfMyoA•FL-PPS structure (chain A) fits well to the SAXS experimental curve ($\chi^2$ = 3.32).* ***(f)*** *When the motor is bound to MgADP, the experimental curve fits poorly to the theoretical curve from the kinked PfMyoA•FL-PR crystal structure ($\chi^2$ = 302),* ***(g)*** *The SAXS experimental curve fits better to the theoretical curve obtained from a model of the PR state in which no kink occurs at the end of the Converter (open conformation) ($\chi^2$ = 19.2).* ***(h)*** *A fit of the theoretical curve in the PR condition with the software Oligomer[4]. The fit has been performed with the closed PfMyoA•FL-PR structure obtained from the crystal with a kink at the pliant region (interface A) and a computed open PfMyoA•FL-PR structure (interface B from the PfMyoA•FL-PPS structure) (see Figure 1). Calculations predict 98% of the sample in the open conformation ($\chi^2$ = 8.69).*

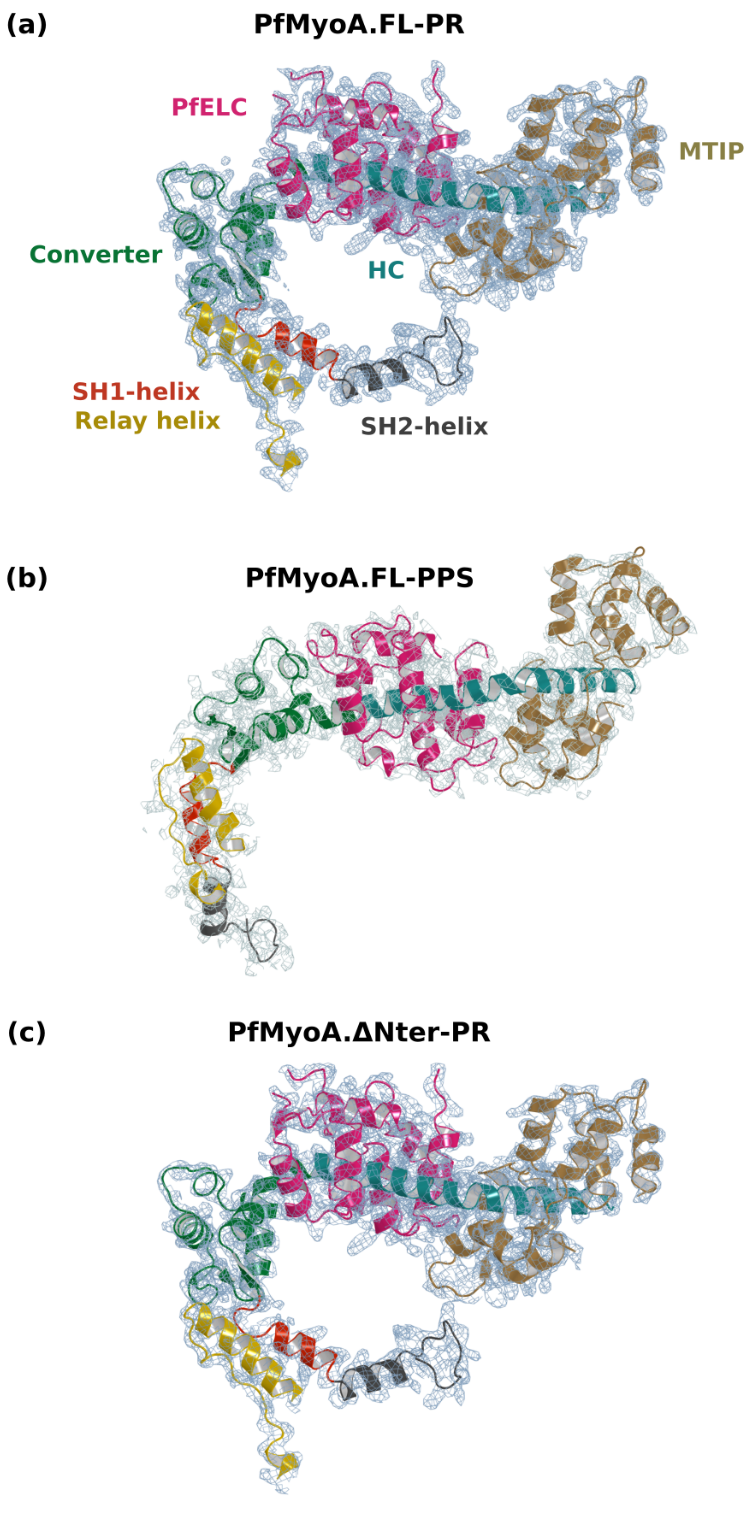

*Supplementary Figure 3 – Electron density in the PfMyoA structures.* *For all structures, the lever arm and the connectors are displayed and the 2Fo-Fc map is shown, contoured at 1.0 σ.* ***(a)*** *PfMyoA•FL-PR at 2.5 Å resolution.* ***(b)*** *PfMyoA•FL-PPS at 3.9 Å resolution.* ***(c)*** *PfMyoA•ΔNter-PR at 3.3 Å resolution.*

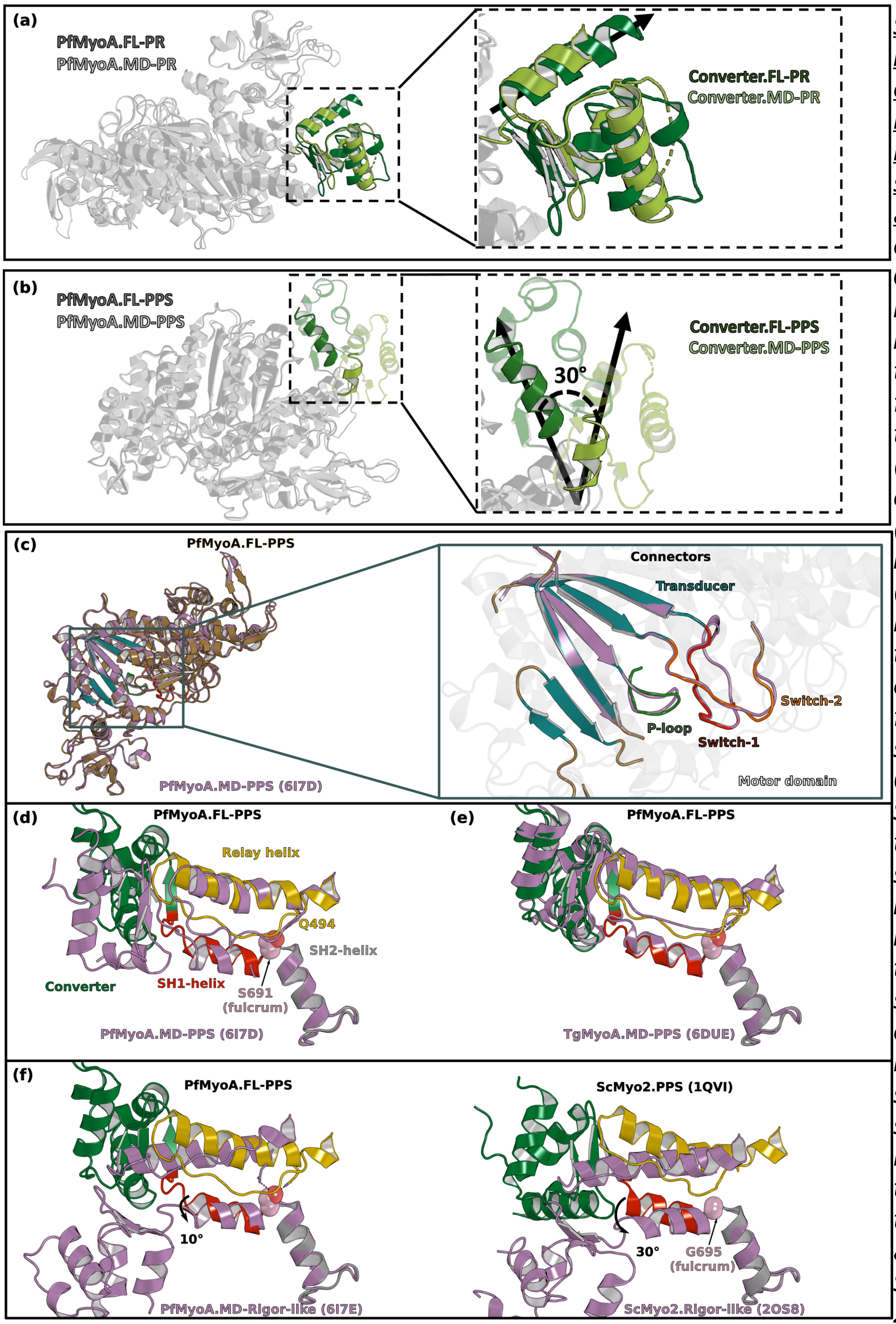

*Supplementary Figure 4 – The orientation of the lever arm of PfMyoA in PPS differs in the structures of the MD and of the FL.* ***(a)*** *The orientation of the Converter in the Post-Rigor state (PR) of PfMyoA is identical in the structures of the full-length (FL) and of the motor domain (MD) constructs (PDB code 6I7D chain A).* ***(b)*** *In the Pre-Powerstroke state (PPS), the lever arm is 30° more primed in the FL structure compared to that found in the MD structure (PDB code 6I7E).* ***(c)*** *Superimposition on the N-terminal subdomain of PfMyoA.FL-PPS and PfMyoA•MD-PPS. On the left, overall view showing that the conformation of the motor domain is similar for the two structures. On the right, zoom on the transducer and on the connectors of the active site (P-loop, Switch-1 and Switch-2) show that the conformation of these elements, which is characteristic of the state, is highly similar. On the following panels, all the structures are superimposed on the SH2-helix.* ***(d)*** *The positions of both the Relay-helix and the Converter (also shown in Supplementary Fig. 6) vary between the FL and the MD constructs. Since the SH1-helix was reported to be immobile in PfMyoA*[5]*, its position only slightly differs between the two constructs.* ***(e)*** *The priming of PfMyoA•FL-PPS is identical to the priming of TgMyoA•MD-PPS (PDB code 6DUE). The communication described between the SH1- and the Relay helices (polar bond between S691 and Q494)*[5] *is similar as well as the orientation of the lever arm.* ***(f)*** *Comparison of the position of the connectors between the PPS and the Rigor-like states in PfMyoA* ***(Left)****, and ScMyo2* ***(Right)****. The priming described in the PfMyoA•FL-PPS structure indicates that the SH1-helix remains mostly immobile: this connector is only rotated 10° during the powerstroke. This rotation is of smaller amplitude compared to that of conventional myosins such as ScMyo2 (~30°).*

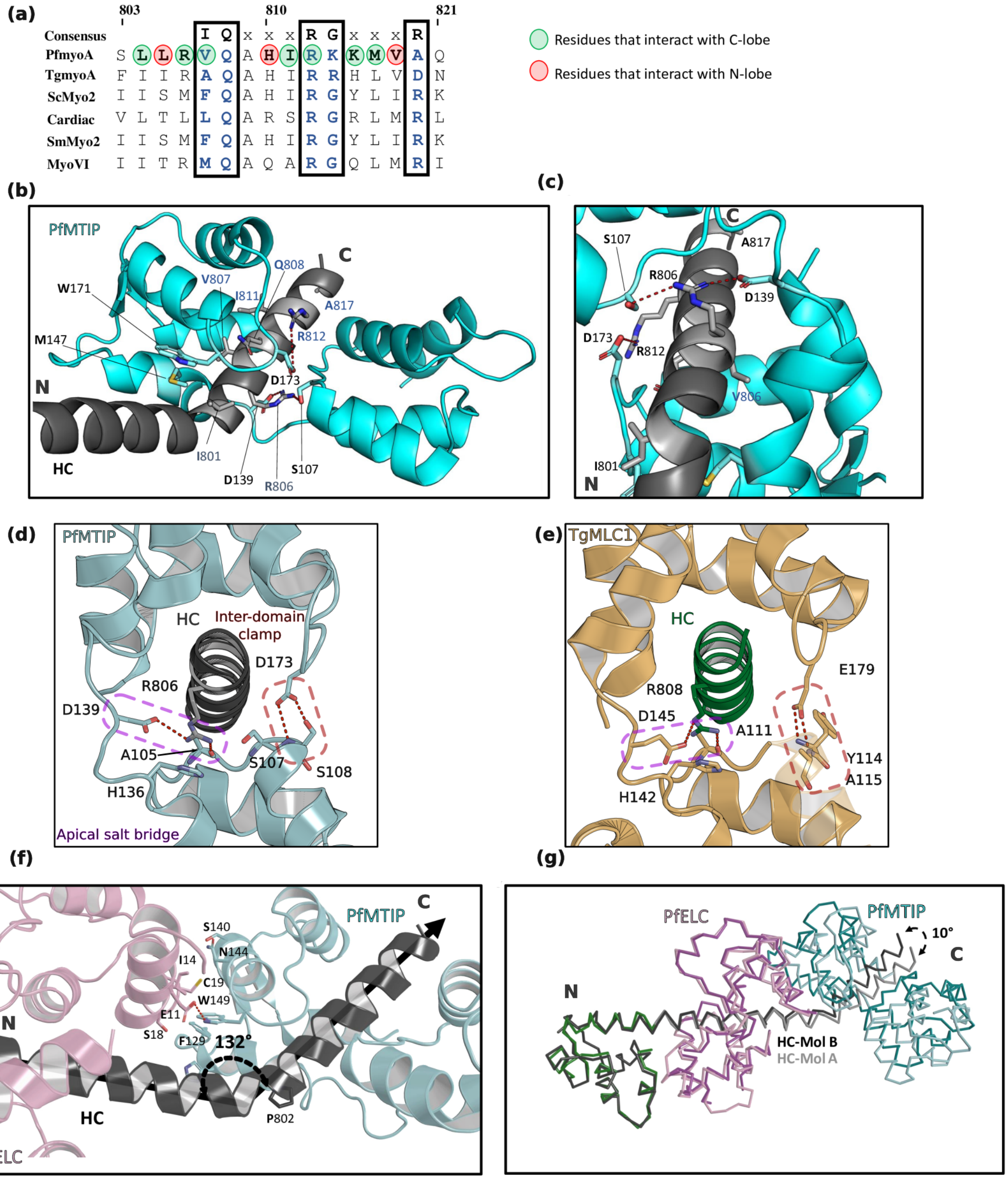

Supplementary Figure 5 – MTIP associated to PfMyoA. ***(a)*** *Sequence alignment of PfMyoA IQ2 with conventional IQ motifs. Residues at consensus positions are colored in blue.* ***(b)*** *MTIP is bound to the heavy chain (HC) in a similar manner as a conventional ELC. Several residues from the consensus are conserved in PfIQ2.* ***(c)*** *Zoom on the specific contacts involved in the MTIP/PfMyoA recognition.* ***(d, e)*** *The recognition of the HC by MTIP is highly similar to the recognition of the HC by TgMLC1 (PDB 5VT9) and the residues involved in the apical salt bridge and the inter-lobe clamp (dotted lines)*[6] are conserved. ***(f) and (g) Interface between PfELC and MTIP.*** ***(f)*** *The interface between PfELC and MTIP is hydrophobic and includes two aromatic residues ($^{PfMTIP}$H150, $^{PfMTIP}$W171) that maintain the relative positioning of the two light-chains. A 132°angle occurs between the two HC helices that contain the PfIQ1 and PfIQ2 motifs. The kink is facilitated by the presence of a proline at the hinge (P802).* ***(g)*** *Alignment of the two molecules of the asymmetric unit in the PfMyoA•FL-PPS crystal structure. The hinge at the PfELC/MTIP interface is a restricted point of compliance since ~10° difference is found for the orientation of the PfIQ2 motif HC helices for these two molecules, when the structures are aligned using the PfIQ1 motif residues.*

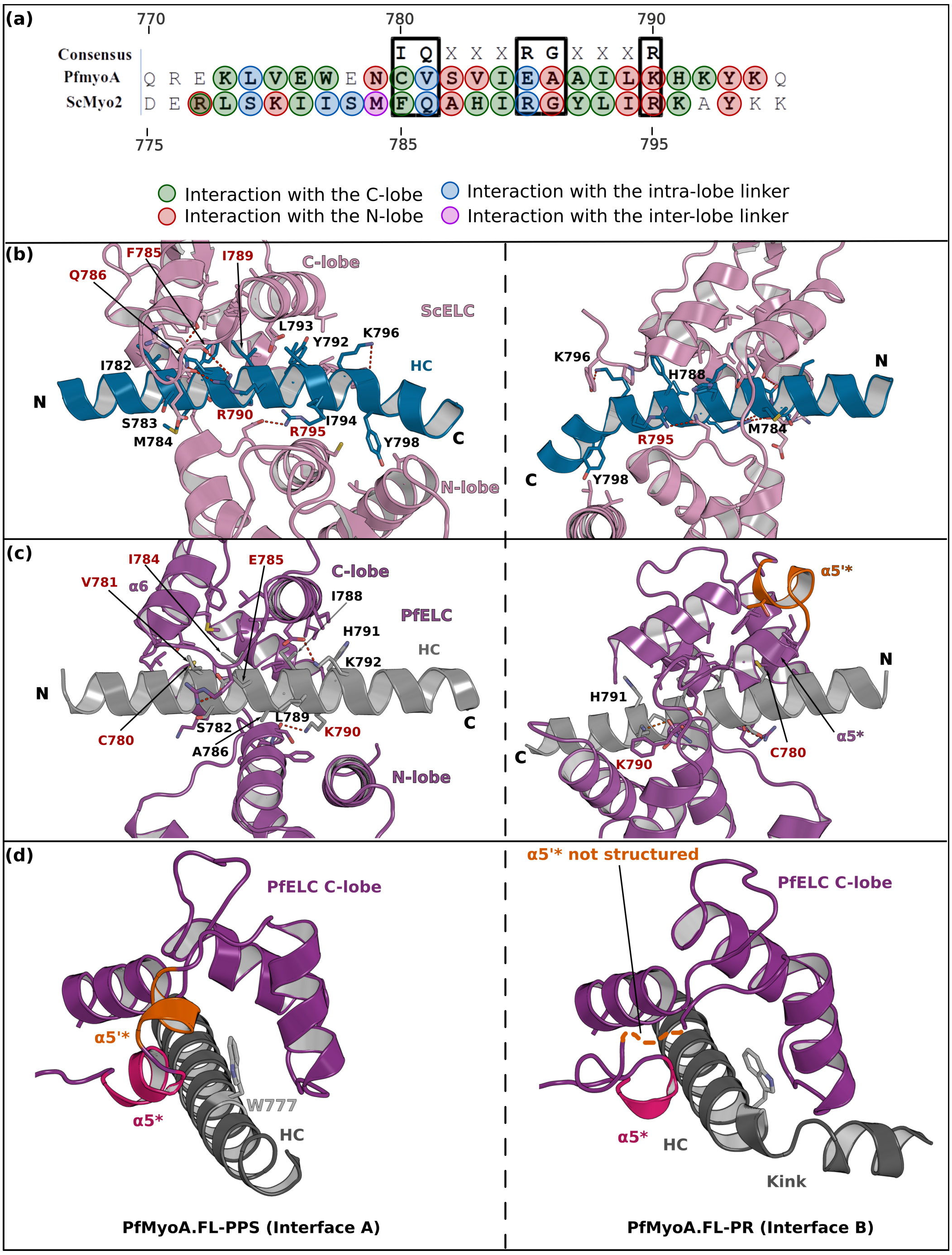

*<u>Supplementary Figure 6 – Interaction between the IQ1 motif and the ELC.</u>* ***(a)*** *Sequence alignment of the IQ1 motif from ScMyo2 and PfMyoA. Residues interacting with the ELCs are contoured following the same color code as defined in* ***Fig. 4****.* ***(b, c)*** *Two different orientations (180° rotation around the y axis) are shown to depict all residues involved in the IQ1 motif/ELC complex recognition, in order to compare* ***(b)*** *ScMyo2 and* ***(c)*** *PfMyoA Residues at consensus positions are labeled in red.* ***<u>(d)</u>*** *<u>The tryptophan W777 stabilizes the α5* and α5'* helices.</u> The left and the right panels show W777 and its environment in the structures of PfMyoA•FL-PPS and PfMyoA•FL-PR, respectively. Note the change in position of the W777 side chain due to its position at the kink of the pliant region. In PfMyoA•FL-PPS, W777 interacts with the α5* and α5'* helices stabilizing their structure. In PfMyoA•FL-PR, the interaction between W777 and the α5* and α5'* helices are lost due to the kink at the pliant region, and a loss of interaction of these helices with the Converter. These helices become disordered.*

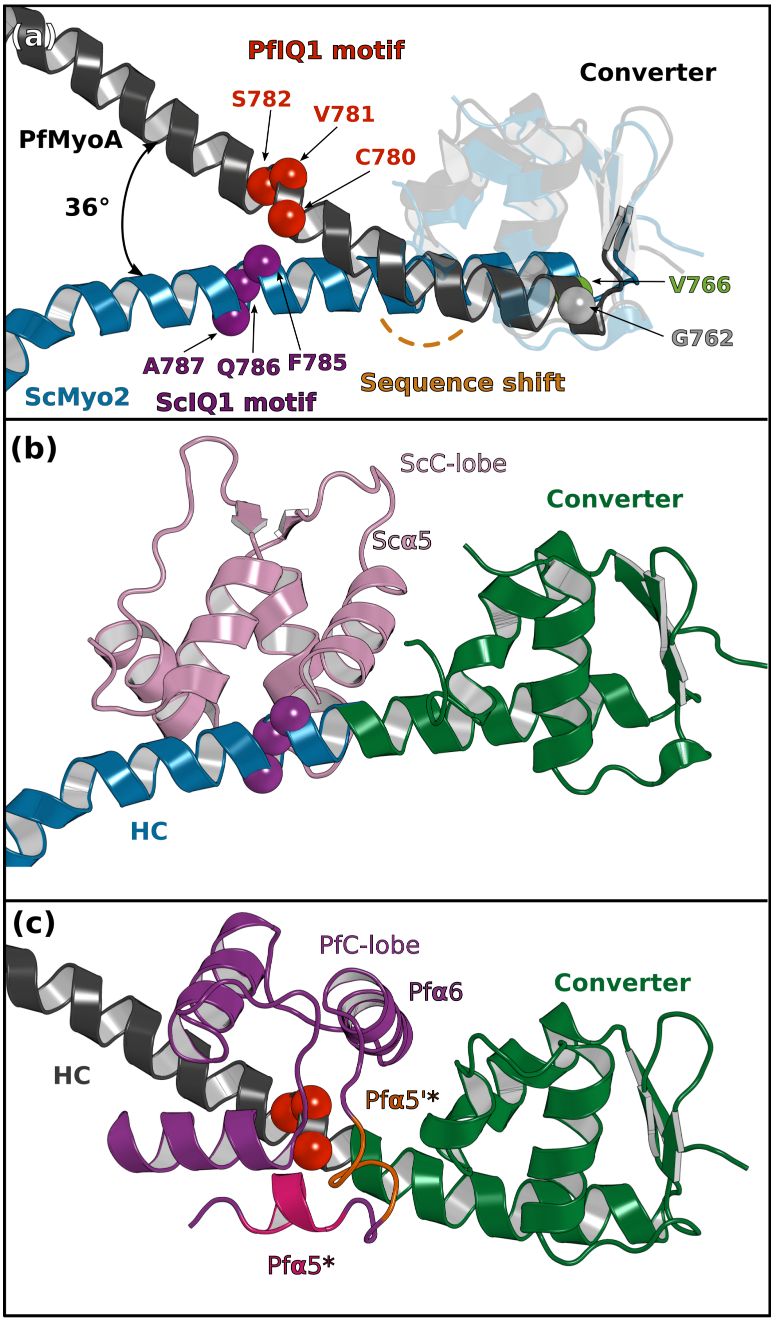

<u>*Supplementary Figure 7 – The Converter/ELC interface differs in PfMyoA and in ScMyo2.*</u> ***(a)*** *The first IQ motif residues of PfMyoA starts one residue downstream to that of ScMyoA (PDB code 1QVI), resulting in a sequence shift in the pliant region and different orientation for the consensus IQ1 motif residues (balls). A difference in the kink at the pliant region accentuates the difference in position of the IQ1 residues in the two lever arms.* ***(b, c)*** *The C-lobe of PfELC and ScELC bound to the IQ1 motif are shown for comparison. The difference in the position of the HC consensus residues result in a different Converter/ELC interface in* ***(b)*** *ScMyo2 compared with* ***(c)*** *PfMyoA (see* ***Fig. 6*** *for details of the interactions).*

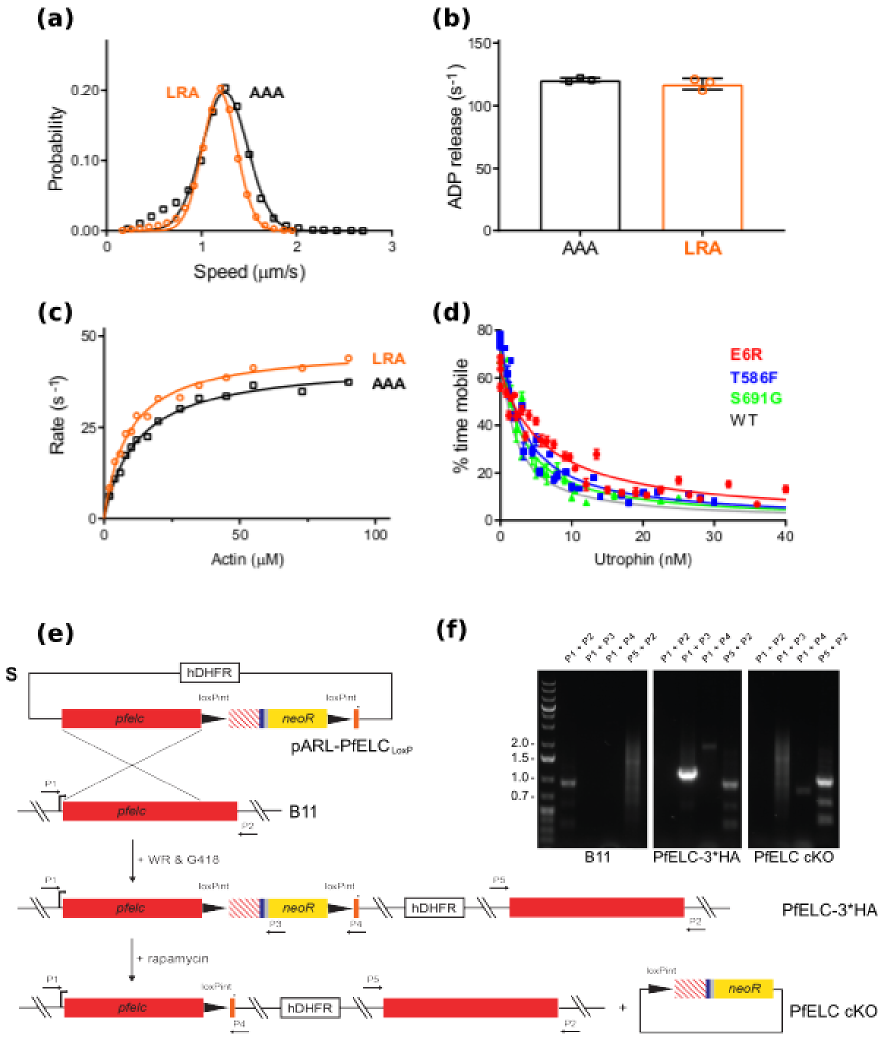

Supplementary Figure 8 – Transient kinetics and genetic engineering of the parasite. **(a)-(d)** Triple mutants that affect the atypical priming of PfMyoA. Speed, ADP release rate, and actin-activated ATPases for LRA (see also data presented in **Fig. 2**) compared with the AAA mutant (LRA: R707L/E711R/Y714A; AAA: R707A/E711A/Y714A), and expanded y-axis for ensemble force measurements (related to main **Fig. 2f**). **(a)** Speed distributions from a representative in vitro motility assay for LRA (1.19 ± 0.18 μm/s, n= 4092) and AAA (1.24 ± 0.24 μm/s, n= 3030). **(b)** ADP release rates from ActoPfMyoA for LRA (117 ± 4 $s^{-1}$) and AAA (122 ± 2 $s^{-1}$). 3 experiments, 1 protein preparation of AAA. **(c)** Actin-activated ATPase activity for LRA ($V_{max}$ = 46.8 ± 1.0 $s^{-1}$; $K_m$ = 9.1 ± 0.6 μM) and AAA ($V_{max}$ = 42.7 ± 0.9 $s^{-1}$; $K_m$ = 12.3 ± 0.8 μM). Error, SE of the fit. 2 experiments, 1 protein preparation of AAA. **(d)**. Ensemble force measurements using a utrophin-based loaded in vitro motility assay, showing more data and an expanded x-axis compared with the main **Fig. 2f**. Temperature, 30°C. **(e)** and **(f)** Genetic integration of LoxP Cre recombinase sites into the Pfelc gene of Plasmodium falciparum. **(e)** Schematics of the targeting plasmid (pARL PfELCloxP), expected selection linked integration (SLI) into the pfelc locus leading to C-terminally tagged PfELC 3xHA and the DiCre-mediated recombinase event resulting in the PfELC conditional KO (cKO). The recodonised version of the pfelc gene is shown as a red striped box, the 3x HA tag is depicted in blue, the T2A skip peptide in gray and the cMyc/Flag tag in orange. The stop codon is indicated by an asterisk and primers used for genotyping are indicated by numbered arrows. **(f)** PCR analysis and confirmation of integration into the pfelc locus in transgenic PfELC 3x HA parasites as evidenced by loss of the wild-type band at ~1 kb (P1 & P2) and amplification of an ~1.2 kb (P1 & P3), ~1.9 kb (P1 & P4) and ~1 kb (P5 & P2) product. Successful rapamycin-induced excision results in the expected loss of the ~1.2 kb band and size reduction of the ~1.9 kb product leading to PfELC cKO.

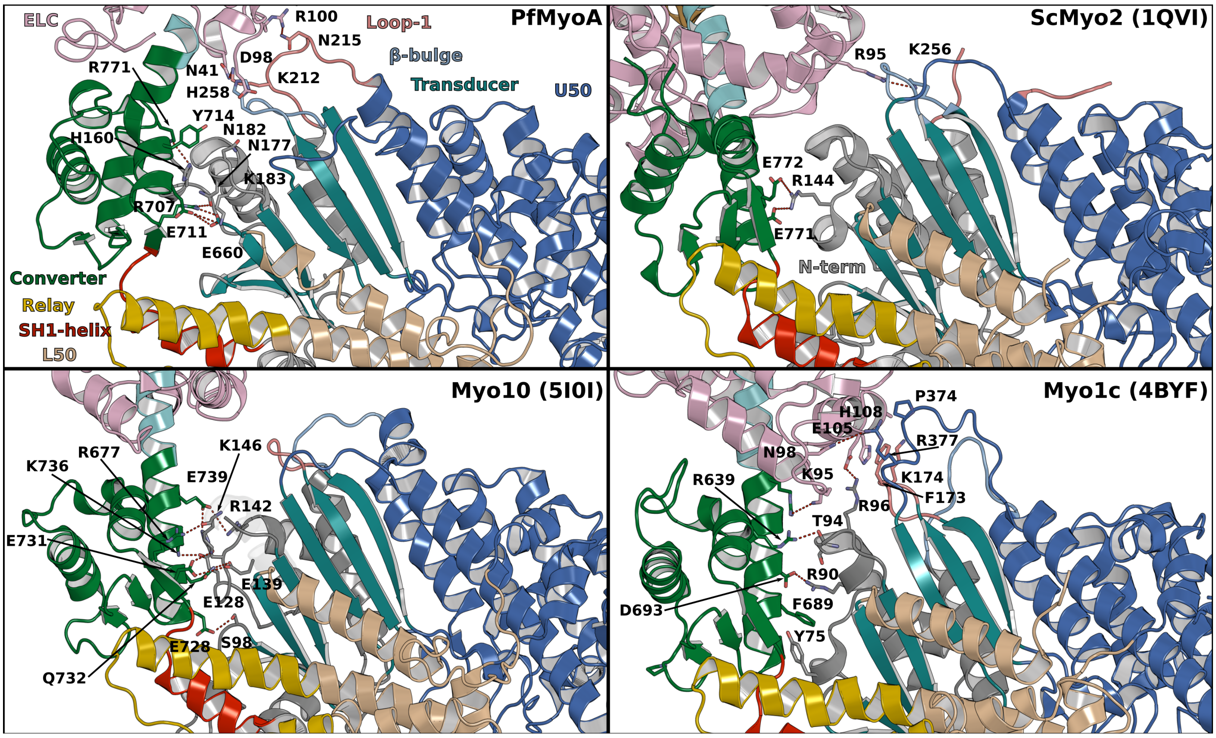

*<u>Supplementary Figure 9 – Interactions between the motor domain (MD) and the lever arm stabilizing the priming in PPS in different myosins</u>. Important interactions occur in PfMyoA between the Converter and the N-terminal subdomain while the PfELC is involved in interactions with Loop1 and the β-bulge of the Transducer. In comparison, limited interactions occur in ScMyo2 consistent with smaller priming of the lever arm. Myo10 is another highly primed myosin but the interactions involve mainly the Converter with the N-terminal subdomain, without contribution from the Transducer. The priming of another highly primed PPS structure from Myo1c requires interaction of the CaM light chain with Loop1 and the HO-linker of the Transducer.*

**Supplementary Data 1: The kink of the lever arm observed in the PfMyoA•FL-PR structure is due to crystal packing**

We investigated by Small-angle X-ray Scattering (SAXS) if the kink in the pliant region observed in the PfMyoA•FL-PR crystal structure was representative of the main conformation adopted in solution. Scattering curves were collected in two conditions, with Mg.ADP (PR condition) or with Mg.ADP.Vanadate (PPS condition). The experimental curves were fitted to theoretical curves computed with crystal structures. In the PPS condition, the experimental curve fits perfectly with the curve from the PfMyoA•FL-PPS structure computed with molecule A ($\chi^2$ = 3.32) (**Supplementary Figure 2e**). In contrast, in the PR condition, there is a poor fit between the curve obtained from the kinked PfMyoA•FL-PR structure and experimental data ($\chi^2$ = 302) (**Supplementary Figure 2f**). We thus modeled an "open conformation" for the PR state without the kink at the pliant region by conserving the Converter orientation and introducing the interface A for interactions between the Converter and the rest of the lever arm (see methods). This model fits better to the data ($\chi^2$= 19.2) (**Supplementary Figure 2g**), indicating that the conformation of the PR in solution is in majority open. To estimate the proportion between open and closed conformations populated in the PR condition, we used the software OLIGOMER[4]. OLIGOMER fits an experimental curve with a multicomponent mixture of proteins and determines the volume fraction of each component. The calculations indicated 98% of the population adopts an open conformation ($\chi^2$ = 8.69) (**Supplementary Figure 2h**). This demonstrates that the closed conformation is due to an artefact of crystal packing which captures a particular orientation of the lever arm region of the PfMyoA•FL-PR structure (**Supplementary Figure 2h**). As previously proposed[7,8], this observation demonstrates that the pliant region displays a controlled dynamics and can be subject of deformation under constraints. This new study adds to the growing list of evidence in this regard, as previously observed with molecular dynamics[9] and by the comparison of crystal structures (PDB code 1BR1[10], PDB code 4BYF[11]). Finally this data indicates that without constraints, the Converter/ELC interface most populated by the PfMyoA FL molecule does not involve a strong kink in the pliant region and is that described as interface A (as found in the PfMyoA•FL-PPS structure) and not that populated in the PfMyoA•FL-PR structure.

**Supplementary Data 2: Recognition of PfIQ2 by MTIP**

The full-length structure also allows to describe the structure of the MTIP bound to its IQ motif in the context of the entire lever arm. Several structures of MTIP bound to its IQ motifs are available from several Plasmodium species[12,13,14] but also from the MLC1 homolog of *Toxoplasma gondii*[6]. MTIP has been crystallized in two conformations, an open-one with a linker helix between the two lobes[12] that was later attributed to have a lower affinity for the IQ motif and a closed conformation that is identical to a canonical ELC conformation and is the form with the highest affinity for the IQ motif [14]. The sequence of the PfIQ2 is conserved among Class XIV myosins and displays several residues from the consensus (**Supplementary Fig. 5a**). In the full-length structure, MTIP is in the closed conformation (rmsd 0.779 Å on 128 atoms with PDB code 4AOM), making similar contacts with the IQ motif as those previously described [12,13,14] (**Supplementary Fig. 5b, 5c**). The polar bond between $^{N\text{-lobe}}$S108 and $^{C\text{-lobe}}$D173 clamping MTIP around the IQ motif (**Supplementary Figure 5b**), the polar interactions between the consensus $^{HC}$R806 and $^{HC}$His810 with the conserved $^{MTIP}$D117 from the inter-lobe linker (**Supplementary Fig. 5c**); as well as the specific contribution of the $^{HC}$K813 with the MTIP C-lobe (**Supplementary Fig. 5c**) are all conserved. Finally, MTIP binds PfIQ2 in a conventional manner, as first described for the ELC binding to IQ motifs of the Myo2 lever arm[15]. The apical salt bridge and the inter-domain clamp[6] are similar for these two apicomplexan MTIPs indicating that the stabilization of MTIP on the myosin HC is similar for these class XIV myosins. Recruitment of MyoA to the IMC via this adaptor light-chain is thus conserved amongst the Apicomplexan (**Supplementary Fig. 5d, 5e**).

**Supplementary Table 1 Data collection and refinement statistics (molecular replacement)**

| | **PfMyoA•FL-PR** | **PfMyoA•FL-PPS** | **PfMyoA•ΔNter-PR** |
|---|---|---|---|
| **Data collection** | | | |
| Space group | $P\ 2_1 2_1 2_1$ | $P\ 2_1 2_1 2$ | $P\ 2_1 2_1 2_1$ |
| Cell dimensions | | | |
| *a*, *b*, *c* (Å) | 89.67 114.69 169.46 | 168.24 287.43 78.61 | 90.08 114.43 170.70 |
| $\alpha, \beta, \gamma$ (°) | 90 90 90 | 90 90 90 | 90 90 90 |
| Resolution (Å) | 25.6-2.55(2.641-2.55)* | 48.4-3.99 (4.133-3.99) | 48.3-3.27 (3.387-3.27) |
| $R_{merge}$ | 0.169 (1.918) | 0.4443 (2.491) | 0.3707 (2.157) |
| $I / \sigma I$ | 10.57 (0.83) | 4.19 (0.64) | 9.02 (1.68) |
| $CC_{1/2}$ | 0.998 (0.505) | 0.989 (0.319) | 0.992 (0.553) |
| Completeness (%) | 99.62 (97.94) | 99.49 (96.58) | 99.88 (99.85) |
| Redundancy | 11.2 (11.5) | 10.3 (10.1) | 12.8 (12.9) |
| **Refinement** | | | |
| Resolution (Å) | 25.57-2.55 (2.62-2.55) | 48.4-3.99 (4.11-3.99) | 48.3-3.27 (3.39-3.27) |
| No. reflections | 641 293 (total) | 343 030 (total) | 358 945 (total) |
| | 57 507 (unique) | 33 243 (unique) | 27 977 |
| $R_{work}$ / $R_{free}$ | 0.199 / 0.248 | 0.237 / 0.273 | 0.184 / 0.232 |
| No. atoms | | | |
| Protein | 8571 | 17 386 | 8490 |
| Ligand/ion | 47 | 66 | 51 |
| Water | 290 | 0 | 4 |
| *B*-factors | | | |
| Protein | 79.82 | 70.84 | 92.81 |
| Ligand/ion | 83.16 | 23.55 | 78.44 |
| Water | 69.70 | | 63.85 |
| R.m.s. deviations | | | |
| Bond lengths (Å) | 0.014 | 0.014 | 0.015 |
| Bond angles (°) | 1.82 | 1.65 | 1.82 |

*Number of crystals for each structure should be noted in footnote. *Values in parentheses are for highest-resolution shell.

**Supplementary Table 2. Kinetic and motility parameters of PfMyoA mutants**

| PfMyoA construct | In vitro motility speed (µm/s ± SD)* | ADP release rate ($s^{-1}$ ± SD)* | ATPase $V_{max}$ ($s^{-1}$ ± SE)* | ATPase $K_m$ (µM ± SE)* | Rate of acto-PfMyoA dissociation by MgATP ($s^{-1}$)[4*] | Basal ATPase ($s^{-1}$)* | Ensemble force (nM utrophin ± SE)* |
|---|---|---|---|---|---|---|---|
| WT[1] | 3.88 ± 0.54 | 334 ± 36 | 138 ± 4 | 30.3 ± 2.3 | 326 ± 9 | 0.3 | 1.40 ± 0.08 |
| | (1) | (1) | (1) | (1) | (1) | (1) | (1) |
| E6R | 1.91 ± 0.35 | 157 ± 8 | 61.8 ± 0.9 | 4.1 ± 0.3 | 857 ± 22 | 0.5 | 4.02 ± 0.31 |
| | (0.49) | (0.47) | (0.45) | (7.39) | (2.62) | (1.67) | (2.9) |
| S691G | 5.06 ± 0.58 | 580 ± 31 | 70.3 ± 1.4 | 15.4 ± 1.0 | 348 ± 7 | 2.7 | 1.99 ± 0.19 |
| | (1.30) | (1.74) | (0.51) | (1.97) | (1.07) | (9) | (1.4) |
| T586F | 4.04 ± 0.44 | 323 ± 54 | 50.1 ± 1.4 | 13.6 ± 1.3 | 471 ± 27 | 2.1 | 2.38 ± 0.18 |
| | (1.04) | (0.97) | (0.36) | (2.23) | (1.44) | (7) | (1.7) |
| LRA[2] | 1.19 ± 0.18 | 117 ± 4 | 46.8 ± 1.0 | 9.1 ± 0.6 | 472 ± 35 | 0.7 | n.d. |
| | (0.31) | (0.35) | (0.34) | (3.33) | (1.45) | (2.3) | |
| AAA[3] | 1.24 ± 0.24 | 121 ± 2 | 42.7 ± 0.9 | 12.3 ± 0.8 | n.d. | 0.5 | n.d. |
| | (0.32) | (0.36) | (0.31) | (2.46) | | (1.67) | |

*Values inside parentheses are normalized relative to WT as 1.

[1] Data from[5]

[2] LRA: R707L/E711R/Y714A

[3]AAA: R707A/E711A/Y714A

[4]Temperature, 20°C. See **Supplementary Table 3** for additional values.

n.d., not determined

**Supplementary Table 3. Dissociation of acto-PfMyoA by MgATP at 20°C.**

| PfMyoA construct | $V_{max}$ ($s^{-1}$) | $K_m$ (μM) |
|---|---|---|
| WT | 326 ± 9 | 240 ± 18 |
| E6R | 857 ± 52 | 351 ± 51 |
| K764E | 801 ± 54 | 257 ± 38 |
| ΔN | 616 ± 45 | 103 ± 25 |
| S19A | 592 ± 40 | 227 ± 44 |
| S691G | 348 ± 7 | 201 ± 12 |
| T586F | 471 ± 27 | 364 ± 50 |
| LRA | 472 ± 35 | 198 ± 42 |

Fits to Michaelis-Menten equation ± SE of the fit.
